# Robustness of a connectome model of the fly walking rhythm generator to modelling choices

**DOI:** 10.64898/2026.11.11.765831

**Authors:** Yue Ju, Yifan Wu

## Abstract

Connectome-constrained models are increasingly used to attribute behaviour to wiring, but their conclusions rest on modelling choices that the connectome does not fix. We independently replicated the rate model of the *Drosophila* nerve cord in which descending neuron DNg100 drives a walking rhythm generator (Pugliese et al., 2026), and asked which of its conclusions survive defensible alternatives to eleven such choices. Changed one at a time, 19 of 20 alternatives left the rhythm intact, including transfer of the wiring to a second connectome. Changed jointly, the rhythm survived in only 20 of 48 model variants when the descending drive was set by rules that do not look at the rhythm. A post hoc sweep of the drive separated two causes. In 12 of the 28 failures a rhythm existed at another drive, in 11 of them a higher one: the published drive lies at the lower edge of the rhythmic range. The other failures concentrated where larger networks met two- or threefold heterogeneity of intrinsic parameters, an interaction that one-at-a-time tests cannot show. In the 31 variants with a rhythm at a common drive of the sweep, the two excitatory interneurons remained necessary (30 of 31) and the inhibitory interneuron dispensable (29 of 31). The rhythm stayed rare among degree-preserving rewired networks when their drive was swept as well, and left–right coupling in the model was weak. The operating point is a modelling choice in its own right, and what a connectome model “produces” should be reported across it.

## 1 Introduction

Synapse-resolution reconstructions now span the adult fly brain and both male and female ventral nerve cords (Dorkenwald et al., 2024; Schlegel et al., 2024; Takemura et al., 2024; Marin et al., 2024; Azevedo et al., 2024; Cheong et al., 2026). They make it possible to constrain dynamical models by measured connectivity rather than by a hand-designed circuit diagram. Connectome-constrained models already predict activity in the visual system (Lappalainen et al., 2024) and reproduce aspects of whole-brain sensorimotor processing (Shiu et al., 2024). At the same time, embodied simulators provide increasingly detailed fly bodies (Lobato-Rios et al., 2022; Wang-Chen et al., 2024; Vaxenburg et al., 2025) and learned whole-body controllers (Aldarondo et al., 2024), and a connectomic graph model trained with reinforcement learning has been reported to control a simulated fly body (Jin et al., 2026). Behaviour alone cannot validate such a model: a deliberately mismatched worm connectome can be trained to produce realistic fly walking (Brunton et al., 2026).

What a connectome model says about a circuit also depends on choices the wiring does not fix. These include which neurons form the network, which synapses count, how transmitter identity sets sign, how single-neuron parameters scale with cell size, how strongly synapses couple, and at which operating point the network is read out. Conclusions of connectome-constrained models can change with training runs (Karaneen et al., 2026), with the connectome features the model is given (Lappalainen et al., 2024), with unconstrained single-neuron parameters (Beiran and Litwin-Kumar, 2025), with the reconstruction of the connectome that the model is built on (Yad and Nabila, 2026), and with the comparison model and the trained solution (Dhawan, 2026); the classic observation that similar rhythms arise from very different parameters (Prinz et al., 2004) makes the same point for pattern generators. Multiverse and specification-curve analyses, developed in psychology and since applied to neuroimaging, make such dependence explicit by running every defensible combination of analysis choices and reporting how often a conclusion survives (Steegen et al., 2016; Simonsohn et al., 2020; Botvinik-Nezer et al., 2020).

We apply this logic to the connectome-constrained rate model of Pugliese et al. (2026), which identified a small central pattern generator downstream of DNg100. The published study reproduces the rhythm in four connectome datasets: the male adult nerve cord (MANC), the female adult nerve cord (FANC), the male central nervous system (male CNS; ventral-nerve-cord synapses only), and the brain and nerve cord (BANC). In the preprint version (Pugliese et al., 2025, Fig. 3h–j) the authors reported that two excitatory interneurons, E1 and E2, were necessary for the rhythm, that the inhibitory interneuron I1 was not, that the minimal circuit of DNg100, E1, E2 and I1 was sufficient, and that rhythm frequency depended on drive. Their model produced a single-leg rhythm, and the published study reports no left–right or inter-leg phase coupling. The authors also tested several choices one at a time: dataset, size normalization, intrinsic heterogeneity and transmitter-specific synaptic strength.

To our knowledge, no previous study has applied this approach—a pre-specified combinatorial design over modelling choices, with a decision rule for each conclusion—to a connectome-constrained circuit model. We report four results (Fig. 1). First, we replicated the rhythm generator and its knockout profile in MANC with an independent implementation. Second, in a combinatorial robustness analysis over eleven modelling choices, the model’s conclusions held under nearly every single alternative, but when choices were changed jointly and the descending drive was assigned by a rule, the rhythm survived in fewer than half of the model variants. Third, a post hoc sweep of the drive showed where that fragility lies: in part the rhythm was present at a drive the rule had not found, and the rest was concentrated where larger networks met high heterogeneity of intrinsic parameters. Fourth, the rhythm was rare among rewired null networks, both at the drive assigned by a rule and across drives, and left–right coupling in the model was weak. The robustness analysis reported in the main text is the second we ran; the first contained alternatives that changed the model by accident, and it is reported with its corrections in the Supplementary Information.

**Figure 1:**
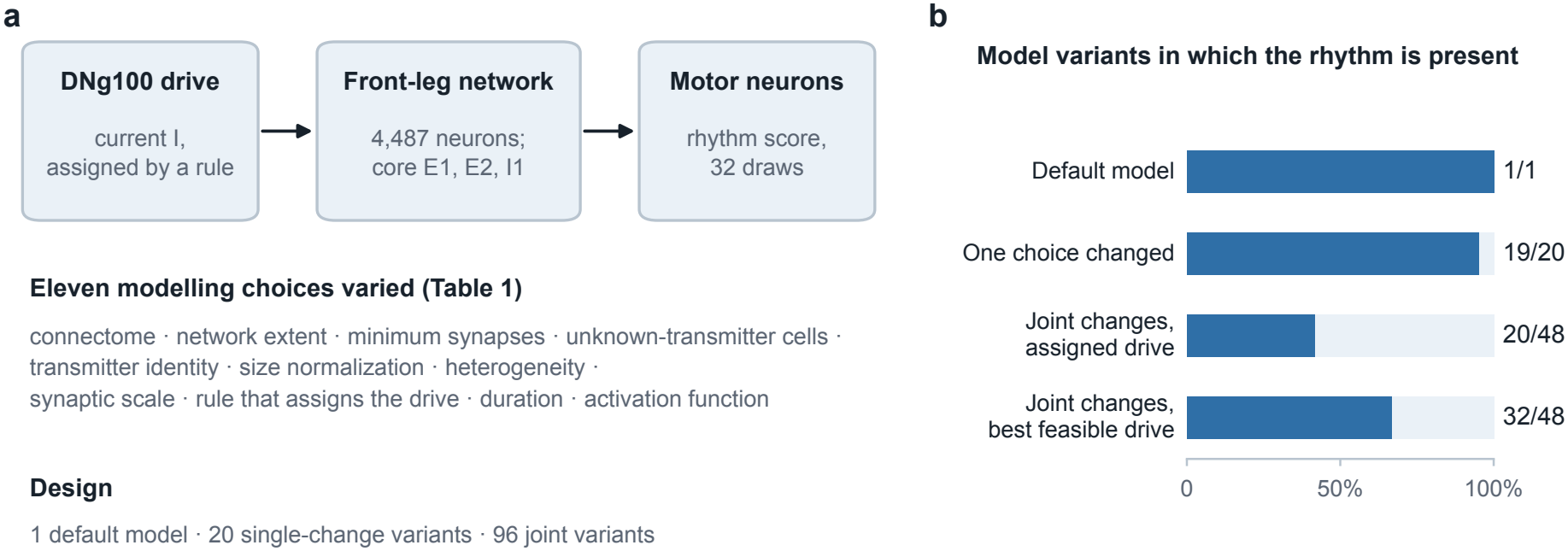
Design and main result. (a) DNg100 is driven by a current assigned by a rule; the rate model of the front-leg network is read out at its motor neurons; eleven modelling choices are varied. (b) Number of model variants in which the rhythm is present: in the default model, with one choice changed, with choices changed jointly at the drive assigned by each variant’s rule, and with choices changed jointly at the best feasible drive of a post hoc sweep. A feasible drive recruits 5–500 neurons; the last bar is the sweep’s primary measure, which also counts variants in which the rhythm was present at the assigned drive. Joint counts are over the 48 variants that keep size normalization.

## 2 Results

### 2.1 Independent replication of the DNg100 rhythm generator and its knockout profile

This section is an independent replication of the knockout and drive results of Pugliese et al. (2026), shown in Fig. 3h–j of their preprint (Pugliese et al., 2025). We reproduced all four results with our own implementation, and also reproduced their finding that bilateral DNg100 activation does not lock left and right legs. Our replication used MANC version 1.0 only; the published study also reproduces the rhythm in FANC, male CNS, and BANC. Three elements are our own extensions: the I1-knockout frequency shift, the linearization of the minimal circuit, and the panel of 64 parameter draws.

We reconstructed the MANC front-leg network from the version-1.0 export and used the authors’ model equation. Each simulation uses one draw of the intrinsic parameters of every neuron from the authors’ distributions. At input current *I* = 250, all 64 parameter draws exceeded the rhythmicity threshold; the mean score was 0.957, the mean dominant frequency was 10.49 Hz, and only 1.94 front-leg motor neurons were recruited per draw (Fig. 2a). The result therefore reproduces a narrow modelled motor rhythm, not broad motor-pool recruitment. Across conditions with rhythmic motor neurons, mean frequency increased from 8.36 Hz at *I* = 175 to 13.19 Hz at *I* = 500 (Fig. 2b). Bilateral DNg100 activation did not produce reliable left–right locking: the across-draw phase vector strength was 0.150 and the mean absolute frequency difference was 1.71 Hz.

**Figure 2:**
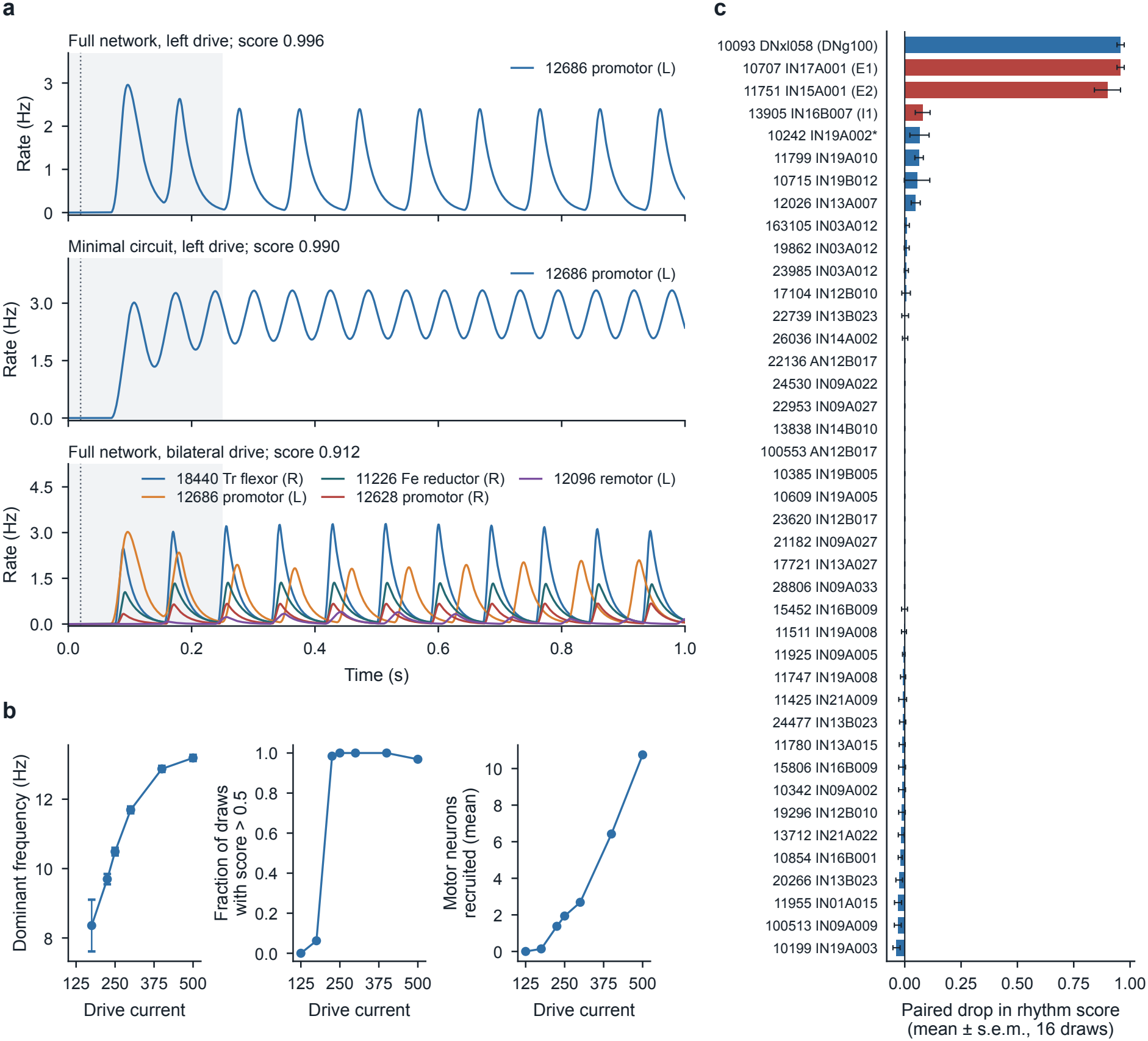
Replication: excitatory-interneuron knockouts suppress a drive-tunable rhythm in the model. (a) Rates of the active front-leg motor neurons in the draw with the median score (draw 44): the full network and the minimal circuit under left DNg100 drive, and the full network under bilateral drive. Shading marks the first 0.25 s, which is excluded from analysis; the dotted line marks drive onset. (b) Dominant frequency (mean ± s.e.m. over draws with rhythmic motor neurons), fraction of draws with a score above 0.5, and mean number of recruited front-leg motor neurons, across drive. (c) Paired drop in the rhythm score when each of 41 cells was silenced; red bars are the core interneurons, and the asterisk marks a cell tested in addition to the 40 most active candidates. Scores average all active front-leg motor neurons; sparse recruitment therefore remains visible beside rhythmicity. Knockout error bars are standard errors across parameter draws and do not represent biological uncertainty.

**Figure 3:**
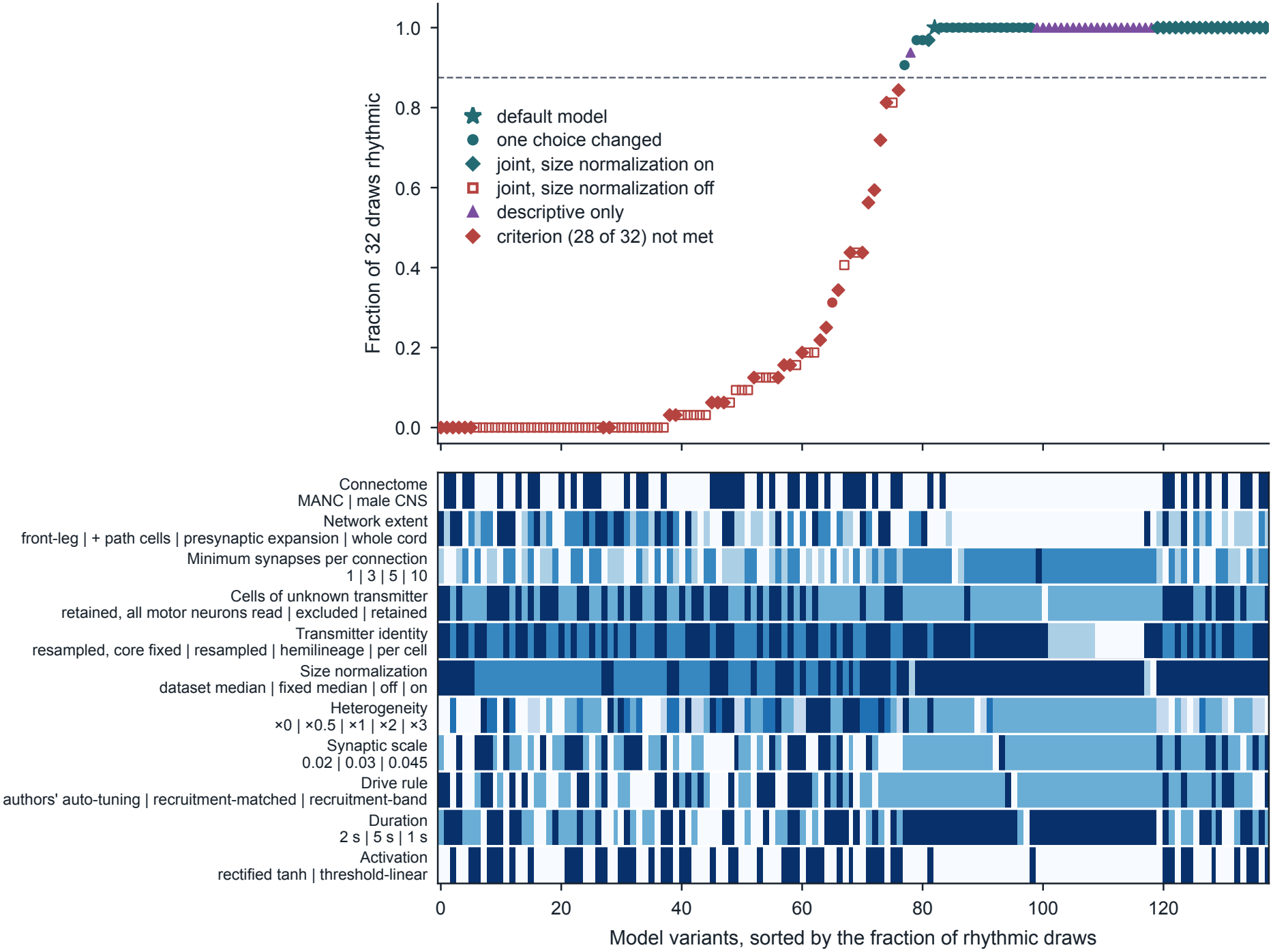
Rhythm generation at the assigned drive in every model variant. Each column is one variant, sorted by the fraction of 32 draws that were rhythmic; the dashed line is the 28/32 criterion. Teal marks variants that met it and red variants that did not. The star is the default model and circles are single-change variants; diamonds are joint variants with size normalization on, on which the plan’s verdict rests; open squares are joint variants without size normalization; purple triangles are descriptive variants, which are excluded from every count. Rows below mark each variant’s level of each modelling choice, from light (first listed) to dark. Variants with threefold heterogeneity use a lower bound on the time constant (Methods).

Knocking out E1 (body ID 10707) abolished the rhythm in all 16 paired draws; knocking out E2 (11751) left one of 16 draws above threshold. Knocking out I1 (13905) left all 16 draws rhythmic; the accompanying 1.15-Hz frequency reduction is our extension (Fig. 2c). The isolated DN–E1–E2–I1 circuit was rhythmic in 98.4% of 64 draws and ran at 16.06 Hz; its approximate linearization contained a damped oscillatory mode at 14.15 Hz. These interventions replicate, within the model, the necessity of E1 and E2 and the dispensability of I1 at this drive and for this panel of parameter draws.

### 2.2 A robustness analysis over eleven modelling choices

#### Conclusions tested

We tested six conclusions of the published model in every model variant, each with a pre-specified decision rule: (i) rhythm generation, DNg100 drive produces a front-leg motor rhythm (at least 28 of 32 parameter draws rhythmic); (ii) necessity of E1 and E2 (silencing either cell abolishes the rhythm in at least 0.875 of the draws that were rhythmic); (iii) dispensability of I1 (silencing it abolishes the rhythm in at most 0.125 of those draws, with a mean score drop below 0.25); (iv) the drive–frequency relation (Spearman *ρ* ≥ 0.9 between drive and mean frequency); (v) specificity to the wiring (the rhythm is rare in degree-preserving rewirings of the variant’s own network); and (vi) absence of left–right locking (at most 0.25 of draws lock after the two sides’ frequencies are matched). Conclusions (ii), (iii), (iv) and (vi) presuppose a rhythm and were evaluated only in variants where (i) held.

#### Choices varied

Table 1 lists the eleven modelling choices we varied. Every alternative had to meet four inclusion criteria, written down before any outcome was scored: it makes one declared change; conventions such as the size median and the motor-neuron readout are computed on the network actually simulated; its value is one the authors used, lies in a range they tested, or rests on a cited measurement; and its drive is set relative to the simulated model. Alternatives that did not meet the criteria were simulated for description only and are excluded from every count below (Methods).

**Table 1:** Modelling choices varied. The default reproduces the replication above. Each alternative met the inclusion criteria.

| Choice | Default | Alternatives |
|---|---|---|
| Connectome | MANC | male-CNS wiring (nerve-cord synapses, MANC-matched cell sizes) |
| Network extent | front-leg network (4,487 cells) | plus the 16 cells on short paths between the left and right cores; plus every cell presynaptic to its premotor cells (17,368 cells); whole nerve cord (22,911 cells) |
| Minimum synapses per connection | 5 | 1; 3 |
| Cells of unknown transmitter | excluded | kept as cells that receive input but have no output, with every traced front-leg motor neuron read out |
| Transmitter identity | per-cell prediction | hemilineage consensus |
| Size normalization | on | off |
| Heterogeneity of intrinsic parameters | ×1 | ×0; ×0.5; ×2; ×3 |
| Global synaptic scale | 0.03 | 0.02; 0.045 |
| Rule that assigns the drive | recruitment-matched | recruitment-band rule; the authors' auto-tuning rule |
| Duration and analysis window | 1 s, window 0.25–1 s | 5 s, window 3–5 s; the authors' 2 s, window 0.25–2 s |
| Activation function | rectified tanh | threshold-linear with a ceiling |

#### Rules that assign the drive

Wiring does not fix how strongly DNg100 is driven, so every variant needs a rule that assigns the current. We used three, none of which looks at the rhythm. The recruitment-matched rule bisects the current, separately for each parameter draw, until the number of active neurons in the front-leg network is within 10% of its value in the default model. The recruitment-band rule, taken from our null-model analysis, starts at 250 and halves or doubles the current until between 5 and 500 neurons are active. The authors’ auto-tuning rule, taken from their code, also starts at 250 and accepts a current at which at least 5 neurons are active, at most 500 are active and at most 100 exceed 100 Hz. We refer to the last two as the band rules, and to the current a rule returns as the assigned drive. In the default model all three return 250.

#### Design

The design has the default model, 20 single-change variants (one for each alternative in Table 1), and 96 joint variants in which every choice except size normalization takes a level drawn at random with each level equally frequent; 48 joint variants keep size normalization on and 48 switch it off. A further 21 variants are descriptive. Each variant simulates 32 parameter draws for conclusions (i)–(iv) and 32 others for (vi); (v) was tested in the default model and the single-change variants only.

#### Status of this analysis

This is the second such analysis we ran on this model. The first was fixed before any variant other than the default was simulated. When we compared it with the authors’ code after reading its outcomes, we found that several of its alternatives had changed the model by accident: brain synapses onto descending neurons in an otherwise nerve-cord network, a size median taken from a different network, a transmitter resampling larger than the classifier’s error, and a drive of 250 carried unchanged into models that need other drives. Two of its apparent findings, loss of the rhythm in male CNS and loss of E1 necessity in larger networks, were consequences of these constructions. The first analysis and its corrections are reported in full in the Supplementary Information (Section S1). The analysis reported here was written to remove those accidents. Its plan was fixed before any of its simulations but after the outcomes of the first had been read, so it is pre-specified but not confirmatory. Its departures from the plan are listed in Methods.

### 2.3 Single changes leave the rhythm intact; at the assigned drive, joint changes do not

#### One change at a time

The rhythm was present in 19 of the 20 single-change variants. All 32 draws were rhythmic when the wiring was transferred to the male CNS, with hemilineage-consensus transmitters, with cells of unknown transmitter retained, with the authors’ two-second simulation and with the authors’ auto-tuning rule, and 29 of 32 were at threefold heterogeneity. The rhythm was lost only when size normalization was switched off (10 of 32), which the authors had reported. With the male-CNS wiring, every conclusion that held in the default model also held: silencing E1 or E2 abolished the rhythm in every draw and silencing I1 in none, the drive–frequency correlation was *ρ* = 0.90, and the rhythm was present in a fraction of draws 0.83 higher than in rewired networks (*p* = 0.03).

The two knockout conclusions failed one at a time only in the two largest networks, and for a reason that is anatomical, not dynamical. Those networks contain the serial homologues of the core cells in the middle- and hind-leg neuromeres, which DNg100 also drives: it makes 185 and 130 synapses onto the left T2 and T3 homologues of E1, against 187 onto the T1 cell. Knocking out the T1 cell alone leaves two driven homologues in place. When the three left E1 homologues were silenced together, the rhythm was abolished in 1.00 (whole cord) and 0.84 (presynaptic expansion) of the draws that had been rhythmic; for E2 the figures were 0.55 and 0.68. Silencing the three left I1 homologues together abolished it in 0.48 and 0.87 of draws, although silencing the T1 cell alone did so in only 0.03 and 0.06. In larger networks the model’s necessity conclusions are therefore statements about a cell type’s serial homologues together, and the dispensability of I1 does not carry over. The drive–frequency relation held in the default model but not in the three larger networks, with a minimum of one synapse per connection, at threefold heterogeneity or at the highest synaptic scale. Specificity to the wiring held in the default model (difference in rhythmic fraction 0.963, *p* = 2*/*99) and failed in the two largest networks, without size normalization, at threefold heterogeneity and at the lowest synaptic scale. The absence of left–right locking failed in the default model itself: 10 of 32 frequency-matched draws locked, against a limit of 8 (see the coupling section below).

#### Joint changes

In the joint design the rhythm was present in 20 of the 48 variants with size normalization on and in none of the 48 without it (Fig. 3; Table 2). The outcome was nearly all or nothing: 20 variants had at least 28 rhythmic draws and 20 had at most 8. Under the plan, a conclusion that holds in fewer than half of the evaluable size-normalized joint variants receives the weakest of four categories of robustness, provided it can be evaluated in at least three quarters of the single-change variants and of those joint variants; otherwise the plan assigns no category. Rhythm generation therefore received the weakest category. Because the other conclusions are evaluated only where a rhythm is present, they could be evaluated in 20 variants against the 36 that the plan required, and received no category. In those 20 variants the two knockout conclusions each held in 19, the absence of locking in 15 and the drive– frequency relation in 5; 15 of the 20 use the two smallest networks, so these counts are not a balanced sample of the design. The plan’s categorical labels for each conclusion are given in the Supplementary Information (Table S1).

**Table 2:** The six conclusions across the robustness analysis. Joint counts are over the 48 joint variants with size normalization on. The knockout, drive–frequency and locking conclusions are evaluated only where a rhythm is present, so their denominators are the variants with a rhythm. The last column is post hoc. For rhythm generation it gives the sweep’s primary measure: a rhythm at some feasible common drive, or a rhythm at the assigned drive (32 of 48). Thirty-one of the 48 had a rhythm at a drive of the sweep; the other variant had a rhythm at its per-draw assigned currents and at no common drive of the sweep. The two knockout conclusions in that column are evaluated at the best drive of those 31, and the drive–frequency relation on the 13 of them with at least four usable drives.

| Conclusion | Fails with these single changes (of 20) | Joint, assigned drive | Joint, drive sweep |
|---|---|---|---|
| Rhythm generation | size normalization off | 20/48 | 32/48 |
| E1 and E2 necessary | presynaptic expansion; whole cord | 19/20 | 30/31 |
| I1 dispensable | presynaptic expansion; whole cord | 19/20 | 29/31 |
| Drive sets frequency | three larger networks; one synapse per connection; heterogeneity ×3; synaptic scale 0.045 | 5/20 | 1/13 |
| Specific to the wiring | presynaptic expansion; whole cord; size normalization off; heterogeneity ×3; synaptic scale 0.02 | not tested | default model only (text) |
| No left–right locking | fails in the default model and with 13 single changes | 15/20 | not tested |

#### The failures sort by the rule that assigned the drive

The proportion of variants with a rhythm differed sharply by the rule that assigned the drive: 12 of 17 variants with the recruitment-matched rule, 6 of 16 with the recruitment-band rule and 2 of 15 with the authors’ auto-tuning rule. The five recruitment-matched failures fell short of 28 of 32 by 1 to 14 draws, with 14, 18, 23, 26 and 27 rhythmic draws. The 23 failures under the two band rules were of a different kind. Both rules accept any current that activates between 5 and 500 neurons, and in 27 of their 31 variants the median assigned drive was the starting value of 250. In 22 of the 23 failures the median number of active neurons in the front-leg network was 30 or fewer, against 84.5 in the default model; in every band-rule variant with a rhythm it was at least 50. A rule that meets the inclusion criteria as written thus behaved, in combination with other changes, like the fixed drive of 250 that the criteria had excluded. The verdict at the assigned drive therefore mixes two questions: whether the model can produce the rhythm, and whether a rule that never looks at the rhythm finds the drive at which it does.

### 2.4 Where the rhythm is lost: an under-driven operating point, then heterogeneity in larger networks

To separate the two questions we swept the drive in a post hoc analysis whose plan was written before its results were read (Methods). Every variant was simulated at 13 common currents in half-octave steps from 62.5 to 4000, with its own simulated duration, and we asked whether at least 28 of 32 draws were rhythmic at some feasible drive, meaning one at which the median number of active neurons lay in the band rules’ own range of 5–500. The plan fixed the interpretation in advance: at least 43 of the 48 size-normalized joint variants would mean that the joint failures are operating-point failures, 36–42 mostly, 24–35 partly. It also recorded our prediction of at least 40.

#### The default drive sits at the lower edge of the rhythmic range

In the default model all 32 draws were rhythmic at 250, 354 and 500, and 3 were at 177; at 707 the network recruited about 1,000 neurons and left the feasible range. The published drive is therefore the lowest drive of the sweep at which the default model is rhythmic. Changes that lower effective excitation, such as a smaller synaptic scale, a larger network whose size median lowers every gain, or the male-CNS wiring for which the authors use a drive of 400, can move the rhythmic range above 250, where the two band rules still find 5 or more active neurons and stop.

#### Twelve of the 28 joint failures were operating-point failures

The plan’s primary measure counts a variant if a rhythm was present at some feasible common drive or at the assigned drive: 32 of the 48 size-normalized joint variants, against 20 at the assigned drive (Fig. 4). Thirty-one had a rhythm at a feasible drive of the sweep; the other one is a presynaptic-expansion variant with a rhythm at its per-draw assigned currents and at no common drive of the sweep. By the plan’s rule the joint fragility is partly an operating-point effect; our prediction of at least 40 was wrong. Of the 28 variants that had failed, 12 were rescued, 11 of them under the two band rules. In 11 of the 12 the lowest rhythmic drive lay above the assigned drive: by one or two half-octave steps in 10 (354 or 500 against 250) and by five in one (1414 against 250). The twelfth, under the recruitment-matched rule, had missed by one draw at a median current of 523 and was rhythmic from 500 upward. At their best drive the variants with a rhythm resembled the default model: a median of 80.5 active neurons in the front-leg network, six recruited motor neurons and 10.6 Hz. The rhythmic range was often narrow, a median of three drives of the sweep and a single one in 9 of 31 variants. An addendum to the plan, written after six variants of the sweep had been read and before any finer simulation, refined the sweep to eighth-octave steps in the 16 variants with a rhythm neither in the sweep nor at the assigned drive. It rescued two, both whole-cord variants with homogeneous parameters and the highest synaptic scale, whose rhythmic range fell between two drives of the coarse sweep. The primary measure rises to 34 of 48 (33 with a rhythm at a common drive) and the interpretation stays the same.

**Figure 4:**
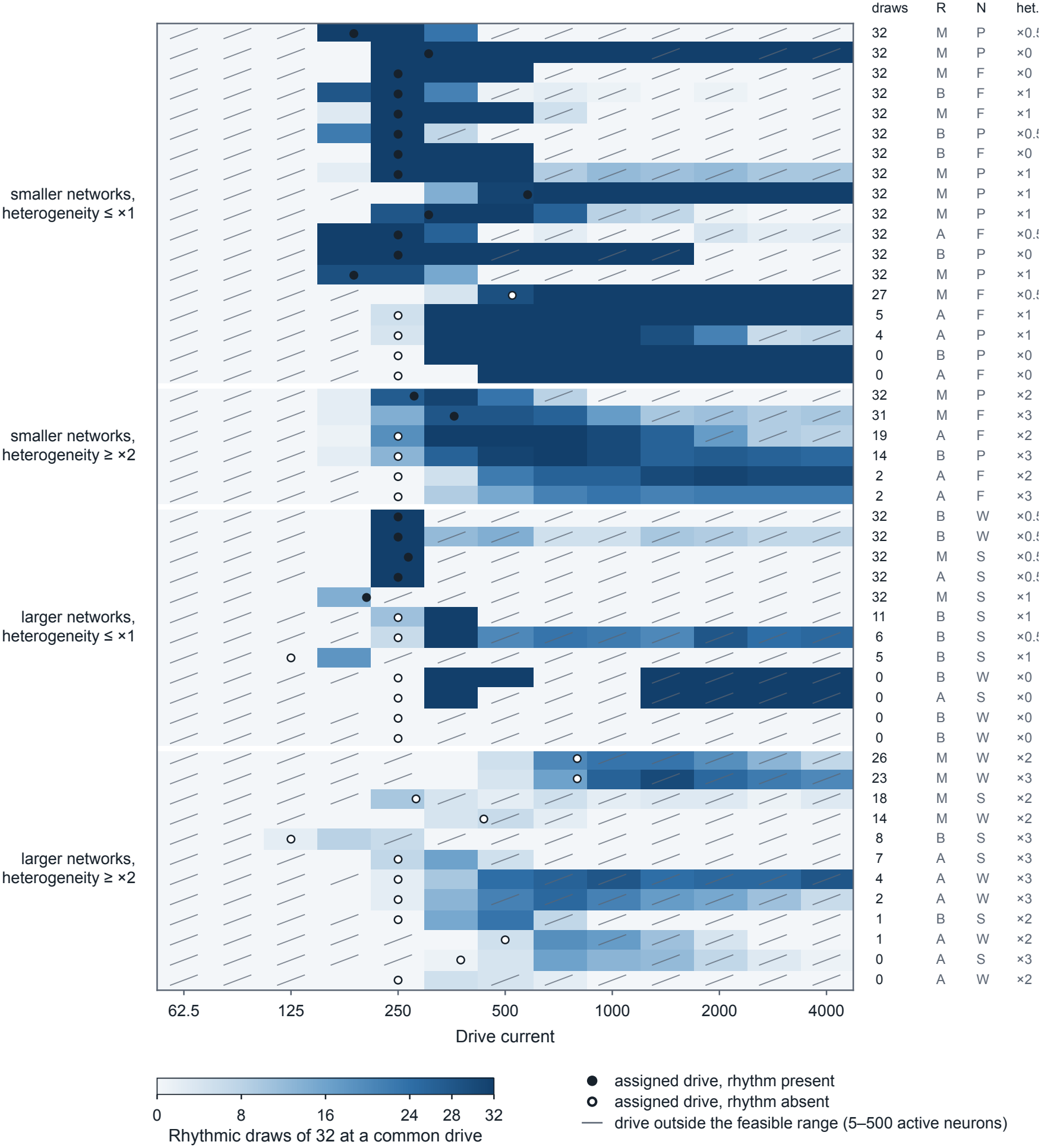
Rhythmic draws across drive in the 48 size-normalized joint variants. Each row is one variant and each column one common drive; colour gives the number of rhythmic draws of 32. A slash marks a drive outside the feasible range of 5–500 active neurons. The circle marks the median drive that the variant’s own rule had assigned, filled if the rhythm was present there; the column headed ‘draws’ gives the rhythmic draws at that drive. Column R gives the drive rule (M recruitment-matched, B recruitment-band, A authors’ auto-tuning), column N the network (F front-leg, P with path cells, S presynaptic expansion, W whole cord), and the last column the heterogeneity. In 6 of the 9 smaller-network rows with an open circle, the circle lies one or two steps below the rhythmic range; of the other three, one has no drive with 28 rhythmic draws, one first reaches 28 at 1414 against an assigned drive of 250, and one has its assigned drive at 523, just above its first rhythmic drive at 500.

#### What the drive does not rescue

The remaining failures were concentrated where two changes met (Table 3). In the two smaller networks with default or lower heterogeneity, all 18 variants had a rhythm, although only 13 had one at the assigned drive: there the drive rule explains every failure. In the two larger networks with two- or threefold heterogeneity, none of 12 did, at any feasible drive. High heterogeneity alone was milder, with 5 of 6 in the smaller networks. Expanding the network, by itself, left a rhythm in most of these variants once the sweep was fine enough: 8 of 12 had a rhythm at a common drive of the coarse sweep and 10 of 12 after it was refined, and one further variant had a rhythm only at its assigned drive, so the primary measure is 9 and then 11 of 12. The feasible range in these networks was a median of two coarse steps wide, because recruitment jumps from tens to thousands of neurons between adjacent drives. Seventeen variants had no rhythm in the coarse sweep. Sixteen of them, those with no rhythm at the assigned drive either, were refined: 13 had heterogeneity of two- or threefold and 15 used a larger network, and the two that the refinement rescued were whole-cord variants with homogeneous parameters. Those 16 were not silent: in the coarse sweep the largest number of rhythmic draws at a feasible drive ranged from 0 to 26, and across all 48 variants 83% of draws were rhythmic at some drive that was feasible for that draw (97–100% in the smaller networks, 75% in the presynaptic expansion and 58% in the whole cord). A common drive is also a stricter demand than the rules make, since they assign the current draw by draw. If each draw is allowed its own drive on the refined sweep, 40 of the 48 variants either reach 28 rhythmic draws or had a rhythm at the assigned drive; the 8 that do neither all have two- or threefold heterogeneity, 7 of them in a larger network, and have 16–27 rhythmic draws. Each of these changes is harmless alone: threefold heterogeneity gave 29 of 32 rhythmic draws and the whole cord 31 of 32 in the single-change variants. The fragility that remains after the drive is accounted for is therefore mainly an interaction between heterogeneity of intrinsic parameters and network size, which no one-at-a-time analysis can show.

**Table 3:**
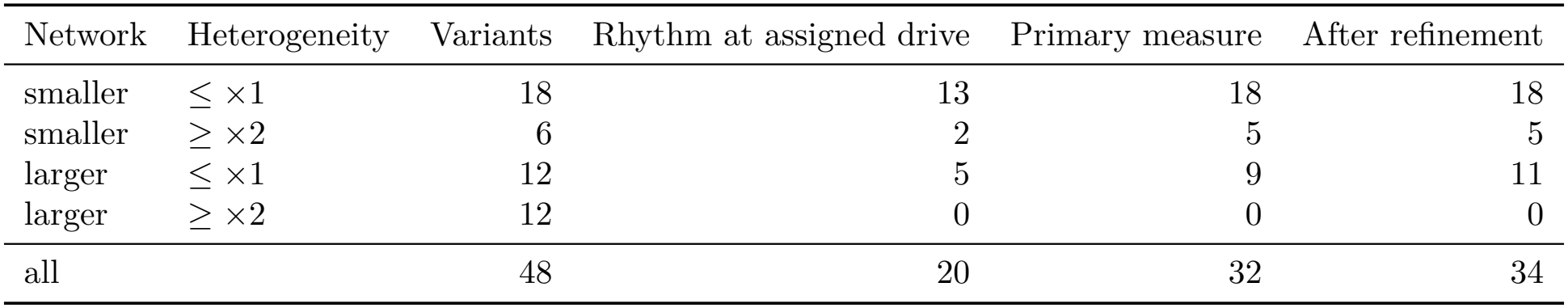
Size-normalized joint variants with a rhythm, by network size and heterogeneity. Smaller networks are the front-leg network and its expansion by 16 path cells; larger networks are the presynaptic expansion and the whole cord. The fourth column is the sweep’s primary measure: a rhythm at some feasible common drive, or at the assigned drive. The last column keeps those variants and adds the ones that gained a rhythm in the eighth-octave sweep. One variant (larger network, heterogeneity ×1) is counted only because it had a rhythm at its assigned drive, so the counts with a rhythm at a common drive are 8 and 10 in its row and 31 and 33 in total; 33 is the count defined in the addendum. The grouping was chosen after the results were seen.

#### Without size normalization

Without size normalization a rhythm was present at a feasible drive of the sweep in 2 of the 48 joint variants, both of them the front-leg network with male-CNS wiring and homogeneous parameters; we had predicted at most 12. This part of the design is poorly resolved even with quarter-octave steps, because recruitment crosses the whole feasible range between adjacent drives: 19 variants had no feasible drive in the sweep and 17 had exactly one (Supplementary Fig. S2). The sweep therefore gives no sign that the dependence on size normalization is an operating-point effect, but it cannot exclude a rhythm in a range narrower than its steps.

#### The knockout conclusions hold in the 31 variants with a rhythm in the sweep; the drive– frequency relation does not extend

At the best drive of each of the 31 size-normalized joint variants with a rhythm in the sweep, silencing the left homologues of E1 and of E2 abolished the rhythm in 30, and silencing the left homologues of I1 left it in place in 29. This replaces the 20 unbalanced variants in which these conclusions could be evaluated at the assigned drive, and includes 8 variants of the two larger networks. In those 8 the three left E1 homologues were necessary in every draw, knocking out the T1 E1 cell alone abolished the rhythm in between 0.28 and 1.00 of draws, and the three left I1 homologues were dispensable (the rhythm was abolished in at most 0.06 of draws), unlike in the single-change variants of the same networks. The exceptions were a male-CNS presynaptic-expansion variant in which silencing the E2 homologues abolished the rhythm in 0.47 of draws, and two male-CNS variants with the lowest synaptic scale in which I1 became partly or fully necessary (0.26 and 1.00 of draws). Drive did not set frequency monotonically over the wider range: 18 of the 31 variants had fewer than four usable drives, and of the other 13 only one reached *ρ* ≥ 0.9, while *ρ* was negative in 11, because frequency levels off or falls at high drive. The published drive–frequency relation holds over the 175–500 range of the default model and should not be extrapolated. Judged by the left T1 E1 cell alone instead of the motor-neuron readout, a rhythm was present in one further variant, so the failures are not failures of the readout.

### 2.5 The rhythm is rare among rewired networks, at the assigned drive and across drives

We asked whether the rhythm of the default model was specific to the measured graph, using four families of null networks: degree-preserving rewiring that keeps the sign of every connection (N1 raw in the figures); the same rewiring rescaled to match the spectral radius of the connectome (N1 scaled); the authors’ permutation of each neuron’s outputs within transmitter and broad cell class (N2); and directed Erdős–Rényi graphs with a fixed number of connections, Dale’s law and the connectome’s weight magnitudes, matched in signed spectral radius (N3). The drive of every network was assigned by the recruitment-band rule, which seeks 5–500 active neurons in the whole network and does not use the rhythm. In a first pre-specified test with 32 networks per family (Fig. 5b) the connectome was rhythmic in all draws with a feasible current, but the design could not reach family-wise significance: the smallest attainable empirical *p* value was 1*/*33 = 0.0303 before correction across 16 tests.

**Figure 5:**
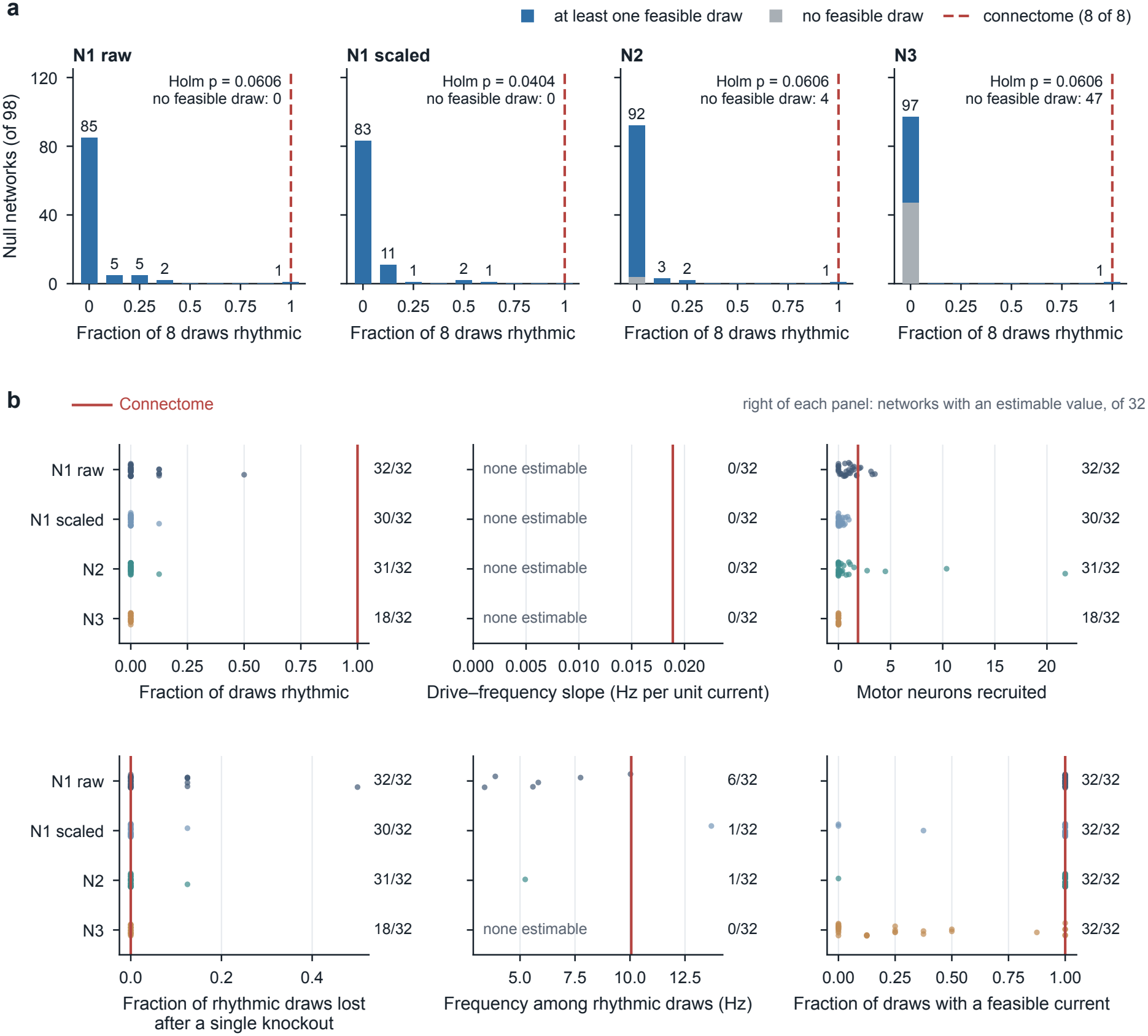
Null networks: the rhythm is rare but not unique. (a) Pre-specified follow-up: fraction of rhythmic draws for 98 networks per family, counting draws without a feasible current as non-rhythmic. (b) Original pre-specified test: properties of the connectome and of the null networks (32 networks per family); the count at the right of each panel is the number of networks in which the property could be estimated. Red marks the connectome; the families differ in how often a feasible current exists and in recruitment. The ensemble that preserves the core circuit is reported in the text and in `d4/report_D4.md`.

A pre-specified follow-up generated 98 networks per family (392 in total; Fig. 5a). Counting draws without a feasible current as non-rhythmic, the mean fraction of rhythmic draws was 0.0370, 0.0332, 0.0191, and 0.0102 for N1 raw, N1 scaled, N2, and N3, and Holm-adjusted *p* values were 0.0606, 0.0404, 0.0606, and 0.0606. Only N1 scaled crossed the family-wise threshold, and that pattern hinged on single ties: in each of the other three families exactly one null network was rhythmic in 8/8 draws and tied the connectome, doubling the unadjusted empirical *p* value from 0.0101 to 0.0202. As an estimate, 0–1 of 98 null networks per family reached 8/8, with Clopper–Pearson 95% upper bounds of 5.6%, 3.7%, 5.6%, and 5.6%. The fractions in the follow-up were higher than those of the original 32-network panel for three families; a post hoc analysis of the stored results attributes this difference to sampling of networks, because rhythmic null draws were concentrated in a few networks.

#### Null networks across drive

The recruitment-band rule stops at the first current that activates between 5 and 500 neurons, and this places the two sides of the comparison at different operating points. The connectome stayed at 250 with 71–94 active neurons, whereas the N1-raw networks, which recruit a median of 2,367 neurons at 250, were halved to a median current of 125 and left with a median of 19 active neurons. In a post hoc analysis we therefore simulated the connectome and the same 98 N1-raw networks at 25 common drives from 44 to 354, and asked of each draw whether it was rhythmic at any drive at which 5–500 neurons were active. The connectome was rhythmic in all eight draws, each at 6–8 of the 25 drives. Among the null networks, 99 of 784 draws (12.6%) could oscillate at some feasible drive, against 29 of 784 (3.7%) at the assigned drive. Thirty-seven of the 98 null networks had at least one such draw, 10 had at least four of eight, and one matched the connectome’s eight. The mean fraction over null networks was 0.126 and the difference from the connectome 0.874 (*p* = 2*/*99 = 0.020), so the rhythm remains rare by the pre-specified rule for specificity to the wiring. Where a null draw could be brought within 50% of the connectome’s recruitment (394 draws), 51 (12.9%) were rhythmic there. The stopping rule thus understated how often the null networks oscillate by about threefold, without changing the conclusion: degree-preserving rewiring removes the rhythm at every feasible drive in most networks, but about one rewired network in ten oscillates in at least half of its draws.

A further pre-specified test asked whether preserving the core circuit is sufficient. We kept fixed the subgraph among both DNg100 cells and the six T1 core cells, allowing no connection among them to be added or removed, and randomized the rest of the graph with swaps that preserve triad counts. The pre-specified test was not supported: the mean fraction of rhythmic draws was 3.44% in 98 such networks against 3.70% in the N1-raw ensemble above (one-sided Mann–Whitney *U* = 4741.5, *p* = 0.60). The pre-specified descriptive thresholds were met, so the plan required that the test stop there; those thresholds do not name which feature of the surrounding network matters, and they do not separate failure of the core circuit from the drive search stopping at the first feasible current. A post hoc follow-up found the left E1 cell rhythmic in 0 of 784 draws of these networks at the assigned drive. Fixing the drive at 250 restored E1 rhythm in 36 of 784 draws, but every one of those draws exceeded the recruitment limit. Within this model, preserving the core circuit was therefore not sufficient for the rhythm in randomized surroundings at the assigned drive; whether the preserved circuit oscillates at some other feasible current was not tested.

### 2.6 Left–right coupling is weak and is not hidden by detuning or by the extent of the network

In the bilateral simulations of the replication, the right front-leg core ran 1.62 Hz faster than the left at equal drive on average (61 of 64 draws had both sides rhythmic). Their short-window phase-locking value (0.33) matched what independent oscillators with the same detunings would give over a 0.75-s window (0.35), so these simulations carry no evidence of coupling (Fig. 6a). Disynaptic and trisynaptic paths between the cores of different legs exist (272 and 36,904 with at least five synapses per connection; there are no direct synapses between cores of different legs), but 15 of the 20 strongest relay cells lie outside the front-leg network.

**Figure 6:**
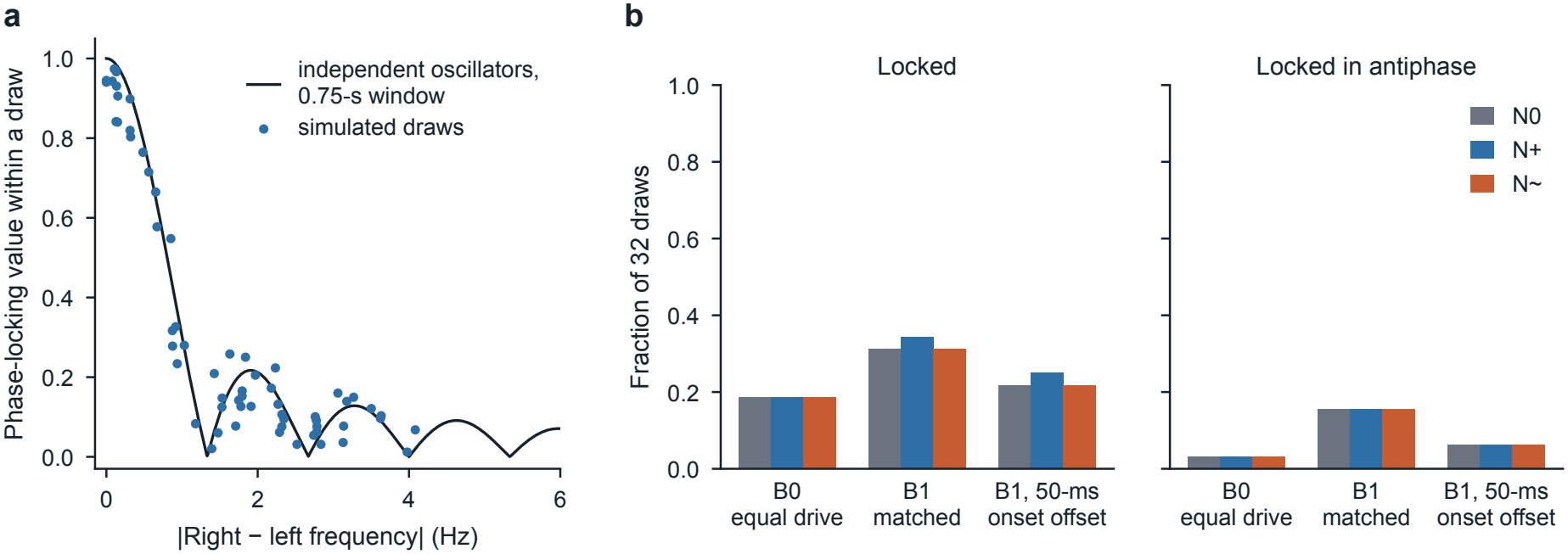
Left–right coupling in the front-leg model. (a) Bilateral simulations of the replication: the short-window phase locking expected from detuning alone matches the simulated values; post hoc analysis of stored results. (b) Fraction of locked draws by network and drive condition. B0 is equal drive, B1 frequency-matched drive, and the third condition adds a 50-ms onset offset to B1; N0 is the front-leg network, N+ its expansion by the 16 path neurons, N~ the control expansion.

A first pre-specified test asked whether detuning hides coupling. At equal drive, 6 of 32 draws locked; after the right drive was adjusted to match the left frequency, 10 of 32 locked. The pre-specified hypothesis that matching increases locking was not supported (Holm *p* = 0.775). The plan’s adjoint method for phase sensitivity failed its validation check, so the direct-pulse method that the plan named as the fallback was used.

A second pre-specified test asked whether the extent of the network hides coupling. Adding the 16 neurons on directed paths of at most three synaptic steps between the T1 cores changed frequency-matched locking from 10 to 11 of 32 draws (Holm *p* = 1), and a control expansion by 16 matched neurons off those paths gave 10 of 32 (Fig. 6b). Post hoc diagnostics showed that the control neurons stayed silent and that only 3–5 of the 16 added path neurons became active. The commissural route proposed by Sapkal et al. (2026) lies at least four synaptic steps from the opposite core and was not tested.

In the robustness analysis, the absence of locking failed in the default model and in 13 of the 20 single-change variants. Within this model, left and right T1 cores lock in a minority of draws, neither detuning nor the tested extent of the network explains the low locking rate, and the tested paths are limited by connections at the five-synapse minimum. This is partly at odds with “no left–right phase coupling” and is better stated as weak coupling. It does not show that biological nerve-cord wiring lacks central coordination: in decapitated flies, left–right alternation persists after proprioceptive removal (Sapkal et al., 2026).

## 3 Discussion

The connectome model of Pugliese et al. (2026) supplied a reproducible rhythm generator whose core conclusions survived nearly every single alternative to its modelling choices, including a second connectome. The combinatorial analysis shows where those conclusions stop holding, and why.

### The operating point is a modelling choice

At the drive assigned by each variant’s rule the rhythm survived in 20 of 48 joint variants, the weakest of the plan’s four categories of robustness. The drive sweep shows that this verdict mixes two things. In 12 of the 28 failures the rhythm was there at another drive, in 10 of them one or two half-octave steps above the drive the rule had assigned. The two band rules that produced most of these failures met our inclusion criteria, because they set the drive relative to the simulated model and never look at the rhythm; in combination with other changes they nonetheless returned the starting current unchanged in 27 of 31 variants. Of their 23 failures, 22 had a median of at most 30 active neurons in the front-leg network, against 84.5 in the default model, whereas every band-rule variant with a rhythm had at least 50. A recruitment range of 5–500 active neurons spans regimes that differ in kind, and the default drive sits at the lower edge of the rhythmic range, so almost any loss of excitation crosses it. Three practical points follow. A rule that assigns the drive should match recruitment to a reference, not merely bound it. A statement that a network does or does not produce a rhythm should be reported over a range of drives, as the authors do when they choose a drive for each dataset. And inclusion criteria applied to how a variant is constructed cannot establish its operating point; recruitment has to be checked on the simulated model before outcomes are read.

### Heterogeneity and network size interact

The failures that the drive did not rescue were not spread over the design. In a grouping chosen after the results were seen, they sat where larger networks met two- or threefold heterogeneity of intrinsic parameters, and in no other group was the rhythm absent in every variant. Each of these changes is harmless when made alone, so a one-at-a-time sensitivity analysis, ours or the authors’, reports the model as robust to both. We did not test a mechanism. The stored recruitment counts show that in larger networks the range of feasible drives is narrow, because recruitment jumps from tens to thousands of neurons between adjacent drives, and a common drive must then suit 32 heterogeneous parameter draws at once. Part of the interaction is thus again an operating-point effect, now at the level of the single draw: with a drive chosen draw by draw on the refined sweep, 5 of the 12 variants that combine a larger network with high heterogeneity reach 28 rhythmic draws, and the 8 variants with a rhythm under none of these criteria all have two- or threefold heterogeneity. Whether intrinsic heterogeneity of this size is realistic for nerve-cord neurons is an empirical question that the model’s robustness now depends on.

### Necessity is a property of serial homologues together

In the 31 joint variants with a rhythm in the sweep, E1 and E2 remained necessary in 30 and I1 dispensable in 29. In networks that include the other leg neuromeres, however, DNg100 drives a homologue of the core in each segment, and the front-leg rhythm survives the loss of a single front-leg cell. The model’s necessity conclusions should therefore be read as statements about the serial homologues of a cell type together. The dispensability of I1 was less stable: its three left homologues together were needed in about half or more of the draws in the single-change presynaptic-expansion and whole-cord variants, but not in the eight joint variants of those networks in which a rhythm was present, and I1 became necessary in two male-CNS variants with the lowest synaptic scale. The drive–frequency relation held over the published range of drives in the default model and did not extend beyond it.

### Specificity to the wiring survives a fairer comparison

The same operating-point problem affected our null-model comparison: the recruitment-band rule placed the connectome at about 84 active neurons and the rewired networks at about 19. Sweeping the drive raised the fraction of null draws that oscillate from 3.7% to 12.6%, and left the connectome’s eight of eight far above the null mean. The rhythm is rare under degree-preserving rewiring, but not as rare as a single stopping rule suggested, and about one rewired network in ten oscillates in at least half of its draws. Two recent preprints bear on this comparison. Liu (2026) shows in other fly circuits that the verdict of a comparison between real and rewired wiring can depend on whether the unmeasured physiology is tuned on the real wiring or on each network separately, which is the distinction between our rule-assigned drive and our sweep. Guan et al. (2026), in a model of the leg motor system with synapse counts as fixed weights, report that rhythm is common among rewired networks and that coordination between antagonistic motor pools is what distinguishes the real wiring. How rare a rhythm is among null networks therefore depends on the model and on its operating point, and should not be read as a property of the wiring alone.

### Left–right coupling

The model’s left and right front-leg cores lock in a minority of draws, and neither frequency detuning nor the tested expansion of the network explains why locking is rare. Longer commissural routes and co-activation of other descending neurons remain untested.

### Two errors of the same kind

This analysis failed in two ways that have the same form. The first robustness analysis contained alternatives that changed the model by accident, which only a line-by-line comparison with the authors’ code revealed; the second removed them and still took its main result from a drive rule that behaved, in combination, like the fixed drive it had excluded. In both, an alternative that is defensible as written was not the same model in the regime where it was run. A robustness analysis is only as informative as the statement that each variant is the same model with one choice changed, and for a dynamical model that statement includes its operating point.

## 4 Methods

File paths in this section are relative to the folder `stage_work/` of the deposit. The stages of the work keep the names they have there: R1 is the replication and its front-leg network; N1, N2 and N3 are the families of null networks and D4 is the test with null networks that preserve the core circuit; D1 and D1b are the coupling tests; M and M2 are the first and second robustness analysis.

### 4.1 Connectome rate model

The network construction began with 144 annotated front-leg motor neurons, all of their presynaptic partners, and descending neurons presynaptic to those partners. We retained traced neurons with known acetylcholine, GABA, or glutamate identity, kept edges with at least five synapses, and removed isolated nodes. The resulting directed graph contained 4,487 neurons and 192,224 edges. Signed weights were stored as *W*_post,pre_, with sign determined by the presynaptic transmitter. For neuron *i*, rate *r*_*i*_ followed

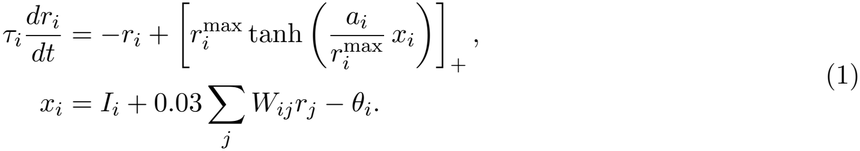

Before scaling, 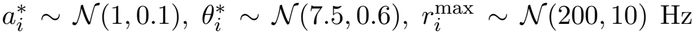, and τ_*i*_ ~ *N*, each truncated to positive values. Gain was divided by relative cell size and threshold was multiplied by it; the size reference was the network median of 676,160,344 units. Simulations used float32 forward Euler integration at 0.2 ms for 1 s from zero rate, saved output every 1 ms, and began drive at 20 ms. The analysis window was 250–1000 ms; an active motor neuron exceeded 0.01 Hz in that window. The rhythmicity score normalized each varying trace, computed a finite-window autocorrelation, required peak prominence of at least 0.05, normalized against sinusoidal references, and averaged over all active front-leg motor neurons; circuit rhythmicity was score *>* 0.5. Dominant frequency was the median autocorrelation frequency among individually rhythmic motor neurons. For three minimal-circuit draws, a high-accuracy DOP853 solution was 0.502 Hz faster on average than the Euler estimate. Differences from the authors’ implementation are listed in `audit/crosscheck_pugliese.md`.

### 4.2 Knockouts

Baseline, drive, minimal-circuit, and bilateral conditions used 64 paired intrinsic-parameter draws. A knockout clamped a target to zero from simulation onset at the same current; loss was the fraction of baseline-rhythmic draws whose score fell to 0.5 or below.

### 4.3 Matched wiring nulls and core-preserving nulls

N1 performed directed double-edge swaps separately by presynaptic sign, preserving signed in/out degrees and each source’s outgoing weight multiset; its raw and signed-spectral-radius-matched forms shared rewires. N2 applied the official shared column permutation within transmitter and broad cell-class groups. N3 sampled a directed fixed-edge-count graph with Dale sign and empirical nonzero magnitudes, then matched the signed spectral radius. For each network and draw, recruitment-only tuning began at *I* = 250 and made at most ten halve, double, or bisection updates to seek 5–500 active whole-network neurons. The follow-up plan targeted 199 networks per family if compute permitted; its pre-declared compute stopping rule ended generation at 98 balanced networks per family. D4 froze the induced subgraph on both DNg100 cells and the six T1 core cells (no edge among them added or removed) and randomized the remaining graph with triad-preserving swaps; 98 networks were compared with the archived N1-raw ensemble by a one-sided Mann–Whitney test.

### 4.4 Robustness analysis (M2)

The remainder of this subsection and Section S1 of the Supplementary Information keep the vocabulary of the plan files, which is used in the repository. M and M2 are the first and second robustness analysis. The plans call the analysis a *multiverse* and a model variant a *universe*, after the statistical literature cited in the Introduction; U0 is the default model and OAT (one at a time) a single-change variant. K1 to K6 are the six conclusions (i) to (vi) of Results, in that order, and the plans call them claims. R1 is the default front-leg network. The same-model test is the set of inclusion criteria. Matched recruitment, the P1 tuner and the authors’ tuner are the recruitment-matched rule, the recruitment-band rule and the authors’ auto-tuning rule. Copies are serial homologues, a clamp silences a cell from onset, loss is the fraction of rhythmic draws in which a knockout abolishes the rhythm, coupling is the global synaptic scale, boundary is the extent of the network, and a label level is a level that met the inclusion criteria. The labels are the plan’s categories of robustness, defined below.

M2 follows`multiverse2/prereg_M2.md.` It was written after M’s outcomes, the cross-check and the post hoc checks had been read, and is not confirmatory. It uses M’s equation, solver, draws, score, readout, rhythm and lock rules and thresholds, with one change: with size normalization on, the size median is that of the cells actually simulated, as in the authors’ code. At the R1 network this is M’s fixed median, so M2’s default variant is M’s. Two departures from the plan are described under Departures below.

#### Same-model test

A level must make one declared change; compute network conventions (size median, readout, sign rule) on the simulated network; rest on a value the authors used, a range they tested, this project’s frozen analyses of the same model, or a cited measurement; and set its operating point by a recruitment target, a recruitment band, or a drive the authors published for exactly that network. Dynamics are never a criterion. The plan records each level’s decision; a construction-only gate confirms that every level matches it.

#### Levels

Label levels, default first: MANC or male-CNS wiring transfer; R1, path, presynaptic or whole-cord boundary; synapse floor 5, 1 or 3; R1 node eligibility or zero-output receivers (traced cells of unknown transmitter kept as receivers, with every traced front-leg motor neuron in the readout); predicted or hemilineage-consensus transmitter (each cell with a hemilineage takes the majority predicted transmitter of its hemilineage across MANC); size normalization on or off; heterogeneity ×1, 0, 0.5, 2 or 3; coupling 0.03, 0.02 or 0.045; matched recruitment, the P1 tuner or the authors’ tuner; 1-s, 5-s or the authors’ 2-s protocol; half-tanh or threshold-linear activation. Descriptive levels, run one at a time and never used for labels, breaking dimensions or predictions: floor 10; receivers plus the non-traced front-leg motor neurons; posterior-calibrated resampling (each cell without a hemilineage changes transmitter with probability min(0.5, 1 − *q*), where *q* is the classifier’s probability of its predicted transmitter), alone and with DNg100’s transmitter fixed; R1’s fixed size median at the presynaptic and whole-cord boundaries; the dataset-wide size median. Not run: a native male-CNS level (no native size data), M’s independent transmitter resampling, and a fixed drive of 250.

#### Variants and claims

The design has the default variant, 20 one-at-a-time label variants, 21 descriptive variants, and 96 joint variants in four balanced cells: size normalization on or off, in two blocks. Claims and thresholds are M’s, with three changes. K2 and K3 clamp the plan’s listed driven-side copies together: the left T1, T2 and T3 copies of E1 (MANC bodyIds 10707, 10559, 10498), E2 (11751, 11767, 12315) and I1 (13905, 13186, 12953), and in male CNS every cell matched to those MANC cells; M’s single knockout is reported beside them. K4 stores recruitment at every drive and flags a drive whose median count of active R1-set neurons exceeds the largest such count at the operating point, without dropping it. K5 uses new nulls in each one-at-a-time variant. The authors’ tuner is their rule, closed in the plan. On the 1-s trial, a current is too strong if more than 500 neurons are ever active or more than 100 exceed 100 Hz, and too weak if fewer than five are active. Too strong halves the current and too weak doubles it until a current on the other side has been seen, after which the current is bisected; there are at most ten iterations, starting from 250.

#### Labels

M’s labels are: Robust, if the claim holds at the default, at every evaluable single-change level and in at least 90% of evaluable joint variants; Interaction-sensitive, if it holds at the default and every single-change level but in fewer than 90% of joint variants; Conditional, if it fails at one or more single-change levels and holds in at least 50% of joint variants; Fragile, if it holds in fewer than 50% of joint variants; and Insufficient, if it is evaluable in fewer than 75% of single-change levels or of joint variants. K5 is labelled from the default and single-change variants only. M’s thresholds apply, with three changes. First, the 50% and 90% joint cuts and the 75% evaluability rule use the joint variants with size normalization on, and the label from all joint variants is printed beside each label. Second, a claim whose only one-at-a-time failure is size normalization off cannot be Robust, and is Conditional when its joint rate with size normalization on is at least 50%. Third, a claim that fails at the default is Insufficient when its joint rate is at least 50% and Fragile otherwise. This denominator was chosen after M’s outcomes were read.

#### T-E2

One network at the whole-cord boundary with size normalization on, three clamp arms at the intact operating current, and two numbers per arm: the loss among baseline-rhythmic draws, and the median frequency of the draws that stay rhythmic. There is no reading rule.

#### Gate and budget

Before any claim variant runs:

- the default variant must reproduce M’s exactly;
- the male-CNS variant with R1’s fixed median must reproduce M’s corrected male-CNS level exactly;
- the expanded boundaries must reproduce the post hoc boundary knockouts;
- every level must match its same-model decision;
- the calibrated resampling’s mean flip rate must lie within 0.02 of the mean of min(0.5, 1 − *q*) over the cells it can flip;
- the consensus assignment’s number of changed cells is compared with a precomputed count (192), and a difference caused by computing the consensus over all of MANC rather than the R1 subset is recorded, not corrected.

A failure of any of the first five, or a consensus difference with any other cause, stops the run. The budget is eight hours on eight workers, projected from timing runs without a safety margin. If the projection exceeds it, the plan cuts in a frozen order: the descriptive variants, then T-E2, then K5 at the whole-cord and presynaptic boundaries and a synapse floor of 1, then the second joint block. The gate passed. The consensus assignment changed 197 cells of the R1 network against the precomputed 192; with the majority taken over the R1 subset it gave 192, so the difference was recorded. The projection was 7.3 h, so nothing was cut.

#### Departures

Both were made before any M2 claim outcome was read and are recorded in `multiverse2/DEVIATIONS.md`. First, the plan defines a copy as a cell of the same type and instance suffix but lists three cells per claim. On MANC v1.0 the definition gives 12 left cells of E2’s type (IN15A001) and 6 of I1’s type (IN16B007). We clamped the lists, which the plan uses in every operative rule, so the expanded-boundary clamps remove 3 of 12 and 3 of 6 left cells of those types. Second, all 19 variants at heterogeneity ×3 produced nonfinite rates. M’s transform floors each parameter at 0.001 times its mean, which lets *τ* reach 0.02 ms, and forward Euler at 0.2 ms diverges when *τ <* 0.1 ms. For *τ* only, the floor at heterogeneity other than ×1 became the 0.2-ms step, which is the limit of the exact update for *τ* much shorter than the step. This overrides the plan’s statement that the far lower tail at ×3 is reported, not corrected. The authors integrate with an adaptive solver and do not face this instability. No value changes at heterogeneity ×2 or below (minimum *τ* 1.40 ms), so the 122 variants that had finished were kept. Labels were computed three ways: with the floor, with every ×3 variant unevaluable, and with every ×3 variant failed under the plan’s rule that a K1 failure makes K2–K4 and K6 unevaluable. They agree for every claim. In the three joint variants that had reached K4 before diverging, the earlier and floored results are identical.

### 4.5 Drive sweeps (post hoc)

Both sweeps were planned after every outcome of the robustness analysis had been read and before their own runs started (`multiverse2/posthoc_drive/plan.md, multiverse2/posthoc_nulldrive/plan.md`). A pilot of the first sweep on two failed variants preceded its plan and is disclosed there. The refinement addendum was written after six variants of the coarse sweep, all whole-cord and two of them size-normalized, had been read. The analysis script was extended after results were seen, with-out changing the measures defined in the plan`(multiverse2/posthoc_drive/NOTES.md)`. Both sweeps reuse the code of the robustness analysis unchanged.

#### Sweep of the model variants

The default model, the 20 single-change variants and the 96 joint variants were simulated with their own duration and analysis window, draws 0–31, with one current for all draws. With size normalization on the currents were 62.5 × 2^*k/*2^ for *k* = 0, …, 12 (62.5 to 4000); without it, 4 × 2^*k/*4^ (4 to 32). A drive is feasible if the median number of active neurons in the whole simulated network lies between 5 and 500, the range of both band rules. A variant has a rhythm in the sweep if at least 28 of 32 draws are rhythmic at some feasible drive. The best drive is the feasible drive with the most rhythmic draws, with ties going to the lowest. At the best drive of every variant with at least 16 rhythmic draws, the listed left serial homologues of E1, of E2 and of I1 were silenced from onset and evaluated by the criteria for necessity and dispensability; the T1 cells were also knocked out singly. The drive–frequency relation was evaluated by Spearman’s *ρ* between drive and mean frequency over feasible drives with at least 16 finite frequencies, requiring at least four such drives and *ρ* ≥ 0.9; variants with fewer than four such drives are reported as not evaluable and are left out of the denominator. The plan’s primary measure is the number of the 48 size-normalized joint variants with a rhythm in the sweep or at the assigned drive, and its interpretation was fixed in advance: the rhythm exists throughout the joint design at 43 or more; mostly at 36–42; partly at 24–35; and the fragility is not explained by the operating point below 24. The plan recorded a prediction of at least 40, and of at most 12 of the 48 variants without size normalization. The addendum’s refined count is the number of variants with at least 28 rhythmic draws at a feasible drive of either sweep (33 of 48); the refined primary measure also keeps the variant with a rhythm only at its assigned drive (34). The per-draw count, which is descriptive, counts a draw if it is rhythmic at some drive of the sweep at which that draw has 5–500 active neurons.

#### Sweep of the null networks

In the default model, the connectome and the 98 stored N1-raw null graphs of the follow-up (regenerated from their seeds, with stored hashes verified) were simulated on the eight draws and 1-s duration of the specificity test (window 0.25–1 s) at 25 common currents, 250 × 2^(*k*−20)*/*8^ for *k* = 0, …, 24 (44.2 to 353.6); the plan writes the first current as 44.2. A draw can oscillate if at some current it has 5–500 active neurons and a rhythmicity score above 0.5. The measure for a network is the fraction of its eight draws that can oscillate, and the rule of the specificity test is applied to it: a difference of at least 0.5 between the connectome and the mean over null networks, with *p* = (1 + *n*)*/*99 *<* 0.05, where *n* is the number of null networks at or above the connectome. The recruitment-matched count takes, for each null draw, the current whose active count is nearest the connectome’s count for that draw at 250, and counts the draw if that count is within 50% and the score exceeds 0.5.

### 4.6 Coupling analyses

D1 used the R1 network with bilateral drive, a 4–30 Hz zero-phase band-pass of each side’s E1 rate, Hilbert phases, and 5-s runs analysed on [3,5] s. A draw was locked if both sides were rhythmic, the phase-locking value was at least 0.9 and the frequency difference was below 0.05 Hz. Matching bisected the right drive to bring the right unilateral frequency within 0.02 Hz of the left. D1b added every traced known-transmitter neuron on a directed path of at most three hops (≥ 5 synapses per hop) between left and right T1 core cells, and a sham expansion matched on transmitter, hemilineage and synapse tercile.

### 4.7 Analysis plans and statistics

The decision rule for each confirmatory test was written down before the data it governs were seen. The replication had no such plan and was not counted as a confirmatory test. Plans written after an outcome had already been read are labelled post hoc wherever they appear. The plan files are included with the code and the summary results. A plan governs only the runs whose outcomes had not yet been seen when it was written. No plan was deposited in an external registry; that each plan preceded its data rests on the project’s own records. Elsewhere in this paper, “pre-specified” refers to these plans.

### 4.8 Independent checks

Each stage was checked by a separate AI-assisted pass that had not written the code under review. These passes read the code and the saved outputs for the R1 network, dynamics, rhythm metric, knockouts and central claims; for the wiring-null stage; for the archive analyses, D4 and its follow-up; for M’s code and results; for M2’s code, its *τ* departure and its results; and for the drive sweeps’ code and results. The check of the drive sweeps compared the code with the plans and recomputed the primary endpoint, the refined count and the boundary-by-heterogeneity table from the stored per-drive counts; it did not recompute the per-draw counts, Spearman’s *ρ* or the null endpoint from the raw scores. The M and M2 results checks recomputed every label, evaluable count and prediction from the claim files. Differences from the authors’ code are listed in `audit/crosscheck_pugliese.md.` A review of the previous version of this manuscript prompted the restructuring around the drive sweeps. Interrupted runs were resumed from the checkpoints already written.

### 4.9 Use of AI tools

Generative AI tools (ChatGPT, Claude and Cursor) were used to assist with coding, analysis and preparation of the manuscript. The authors reviewed all content and take full responsibility for it.

## 5 Limitations

The neural conclusions apply to a rate model with sampled intrinsic properties, not to experimental necessity. The network omits gap junctions, neuromodulation, detailed membrane dynamics, sensory feedback, and uncertainty in reconstruction. Only a narrow subset of motor neurons was recruited, and the numerical solver and simulated duration differed from the authors’. Our MANC release, the cells we admitted to the network and the motor-neuron readout differ from the authors’; the effect is small in the default model but untested in combination. Proportions of joint variants depend on the balanced mix of levels in the design; they are not probabilities that a conclusion is true of the fly. The robustness analysis reported in the main text was specified after the outcomes of the first had been read and is not confirmatory; its denominator (joint variants with size normalization on) was also chosen then, which is why the plan’s labels are given for both denominators in the Supplementary Information. Its lower bound on the time constant at threefold heterogeneity was introduced after the run had started, though before any outcome was read; no verdict depends on it, but one prediction does. In the expanded networks, silencing serial homologues removes one left cell per segment of E2’s and I1’s cell types, not every cell of those types. The knockout, drive–frequency and locking conclusions can be evaluated only where a rhythm is present, at the assigned drive a subset dominated by the two smaller networks. The single-change outcomes of the specificity test near *p* = 0.05, and left–right locking in the default model (10 locked draws against a limit of 8), lie close to their criteria. The coupling analyses tested paths of at most three synaptic steps and unilateral or bilateral DNg100 drive only. The drive sweeps were planned after every outcome of the robustness analysis had been read. They choose the drive by the rhythm, which the inclusion criteria forbid for a verdict, and so answer a different question from the pre-specified decision rules. They use one current for all draws of a variant, half-octave steps, and the band rules’ range of 5–500 active neurons, which was set for networks of about 4,500 cells. The sweep of null networks covers one family, one model and eight draws per network. AI tools were used at every computational stage of this work; the independent checks reduce some implementation risk but do not replace experimental replication or line-by-line human review of the code. The analysis plans were not externally registered.

## Declarations

## Ethics approval

Not applicable. The study involved no human participants or animals; it used published connectome data only.

## Data availability

MANC version 1.0 public exports are available from the public bucket gs://flyem-manc-exports/v1.0; male CNS version 1.0 exports from the public bucket gs://flyem-male-cns. The analysis-plan files and the saved summary results are deposited at Zenodo (https://doi.org/10.5281/zenodo.24126403) under CC BY 4.0.

## Code availability

The code is deposited at Zenodo with the data (https://doi.org/10.5281/zenodo.24126403) under the MIT licence.

## Author contributions

Yue Ju was the main contributor to the study: its design, the analyses and the writing of the manuscript. Yifan Wu wrote code and provided technical support. Both authors revised the manuscript and approved the final version.

## Supplementary Information

### S1 The first robustness analysis (M) and what the cross-check changed

This section reports the first robustness analysis as it was frozen and run, together with the corrections that led to the second (M2), which is the analysis in the main text. M’s labels are frozen and are not replaced by M2’s. This section keeps the vocabulary of the plan files; the glossary at the start of Methods, subsection Robustness analysis (M2), gives the correspondence with the main text.

#### Design

M was specified before any non-default variant was simulated. It scored six claims in every variant: K1, DNg100 drive produces a front-leg rhythm (≥ 28*/*32 rhythmic draws); K2, E1 and E2 are each necessary (loss ≥ 0.875 for both); K3, I1 is not necessary (loss ≤ 0.125, mean score drop *<* 0.25); K4, drive sets frequency (Spearman *ρ* ≥ 0.9 over retained drives); K5, the connectome rhythm is rare in degree-preserving nulls built on each variant’s weights; and K6, left and right T1 rhythms do not lock after frequency matching (locked fraction ≤ 0.25). Ten dimensions were varied: dataset (MANC, male CNS), boundary (R1, path-complete, presynaptic expansion, whole nerve cord), synapse floor (1, 3, 5, 10), transmitter identity (predicted, resampled from classifier probabilities), size normalization (fixed median, recomputed, off), heterogeneity (0, 0.5, 1, 2 × SD), coupling scale (0.02, 0.03, 0.045), operating current (recruitment-matched, P1 tuner, fixed 250), trial window (1 s, 5 s) and activation (half-tanh, threshold-linear). The design ran U0 (all defaults, identical to R1), 26 one-at-a-time (OAT) variants covering 19 levels, 64 balanced joint variants, and one appendix variant: 92 variants in all. Labels were fixed in advance: Robust, Interaction-sensitive, Conditional, Fragile, or Insufficient when a claim is evaluable in fewer than 75% of OAT levels or joint variants.

#### Results as frozen, after the disclosed deviations

U0 reproduced the stored R1 vectors exactly and the archived P1 null result (difference 0.963, *p* = 2*/*99). K1 held at 17 of 19 OAT levels; it failed when transmitter identities were resampled (6 of 8 resamples held; 7 were required) and when size normalization was switched off (10/32 rhythmic). In the joint design K1 held in 21 of 64 variants (32.8%), so its label is Fragile (Fig. S1). Because K1 failed in most joint variants, K2, K3, K4 and K6 were evaluable in only 20–21 of 64 joint variants and are Insufficient. K5 is Conditional: it held at U0 and failed at OAT levels of four dimensions (transmitter resampling, size normalization off, homogeneous intrinsic parameters, and the lowest coupling scale). K6 failed at U0 itself: 10 of 32 frequency-matched draws locked (0.312), in agreement with D1 (main text). Of four frozen predictions, one was true (K5 holds at U0 and fails at OAT levels of at least two dimensions) and three were false.

#### What the cross-check changed

We then compared every M default and alternative with the authors’ code and data (`audit/crosscheck_pugliese.md`, Pugliese_2026 commit 10e7661). The defaults matched in equation, parameter distributions, sign rule, synapse floor, score, active threshold and drive. They differed in dataset release (our MANC v1.0 against v1.2.1), in node eligibility (4,487 against 4,604 cells; 4,486 shared), in readout (107 against 144 front-leg motor neurons), in solver, and in trial length. The cross-check also found alternatives that changed the model by accident; the four that matter for labels are listed here.

1. **Male CNS**. Our male-CNS level used all-ROI weights, so brain synapses onto descending neurons entered a network with no other brain cells. The authors restrict male CNS to synapses in nerve-cord ROIs.
2. **Expanded boundaries**. The presynaptic and whole-cord boundaries used the fixed R1 size median, whereas the authors recompute the median on the simulated network. On our data this made every neuron’s gain 1.31 and 1.39 times higher, with correspondingly lower thresholds. These networks also contain the T2 and T3 copies of the core: DNg100_L sends 185 and 130 synapses to the left T2 and T3 copies of E1, against 187 to T1, so knocking out E1 in T1 alone leaves two driven copies in place.
3. **Transmitter resampling**. Each cell’s transmitter was drawn independently from its classifier probabilities. This flipped about 17% of cells per resample, more than the 6% neuron-level error of a brain-wide transmitter classifier (Eckstein et al., 2024) and against the lineage rule of the nerve cord (Lacin et al., 2019). The K1 failure at this level rests on one resample that turned both E1 and E2 into GABA cells, with per-cell probabilities of 0.009 and 0.017.
4. **Fixed operating current**. A current of 250 was carried unchanged into models whose authors use other drives (400 for male CNS, for example).

#### Post hoc checks, reported as post hoc

Both checks were planned before they were run, but after M’s outcomes had been read. In the male-CNS check, our pipeline reproduced the authors’ result on their nerve-cord network: 32 of 32 draws were rhythmic at their sizes and drive, and our frozen rules on their weights also gave 32 of 32. Replacing only the edges into descending neurons in our network restored the rhythm (32/32). Adding our edges into descending neurons to their network removed it (0/32) and reproduced the recruitment cliff. Edges into descending neurons carried 25.0% of total |*W* | at our level, against 2.7% in MANC. We then restricted the male-CNS level to nerve-cord ROIs (`multiverse/DEVIATIONS.md,` part 5) and reran its 33 variants. That change came after the outcomes were seen and is reported with both versions: K1 held at the corrected level, and its joint rate rose from 8/62 to 21/64, which leaves its label Fragile.

In the boundary check we clamped the same-type copies together. At the whole-cord boundary with the network’s own median, the T1 E1 knockout lost the rhythm in 0.55 of baseline-rhythmic draws, and the three left E1 copies together in 1.00. The presynaptic boundary gave 0.68 and 0.84. M’s “E1 stops being necessary” therefore mostly reflected same-type copies in other segments, and we withdraw it as a result. E2 behaved differently: clamping all left E2 copies lost the rhythm less often than clamping T1 alone (0.55 against 0.65 at the whole cord), and the remaining rhythm was slower (8.2–8.7 against 9.7–11.5 Hz at baseline). This pattern was not predicted and is not interpreted here; M2 reports one pre-specified contrast on it, without a reading rule (Section S2).

#### Methods for M

M used the R1 equation, solver, stored draws, score and readout in every variant (draws 0–31 for K1–K4, 0–7 for K5, 32–63 for K6). Cells outside R1 received bodyId-keyed parameter draws. The default operating current matched each draw’s count of active R1-set neurons to within ±10% of its R1 value by bisection. K4 used drives of 175, 225, 250, 300, 400 and 500 scaled by the operating current, dropping drives with fewer than 16 finite frequencies. K5 used the P1 endpoint and 32 new N1-raw nulls per OAT variant, and was not run in joint variants. K6 matched the right drive to the left frequency and used 5-s bilateral runs analysed on [3,5] s. Joint variants paired levels at random with each level appearing equally often. Departures from the plan are recorded in `multiverse/DEVIATIONS.md`, parts 1–5; parts 4 and 5 were made after outcomes were read and are reported with both versions.

#### M and M2 side by side

Every M2 label equals M’s, for size normalization on and pooled alike (Table S1). The agreement is weak evidence that the labels are stable. The two designs share the default variant and most levels, and the claims whose labels could have moved were already Insufficient for lack of evaluable variants or Conditional with several one-at-a-time failures. Each change in the one-at-a-time failure lists coincides with a changed rule: transmitter resampling left the design; the copy clamp added the whole-cord boundary to K3’s failures; heterogeneity ×3 entered K4’s and K5’s; the network’s own size median at expanded boundaries and new null seeds changed K5’s boundary and homogeneous-parameter outcomes; and K6’s list follows the new level set. At homogeneous intrinsic parameters K5 failed in M (*p* = 0.061) and held in M2 (*p* = 0.030) on the same network with new null seeds, so its single-change outcomes near *p* = 0.05 depend on the null sample.

**Figure S1:**
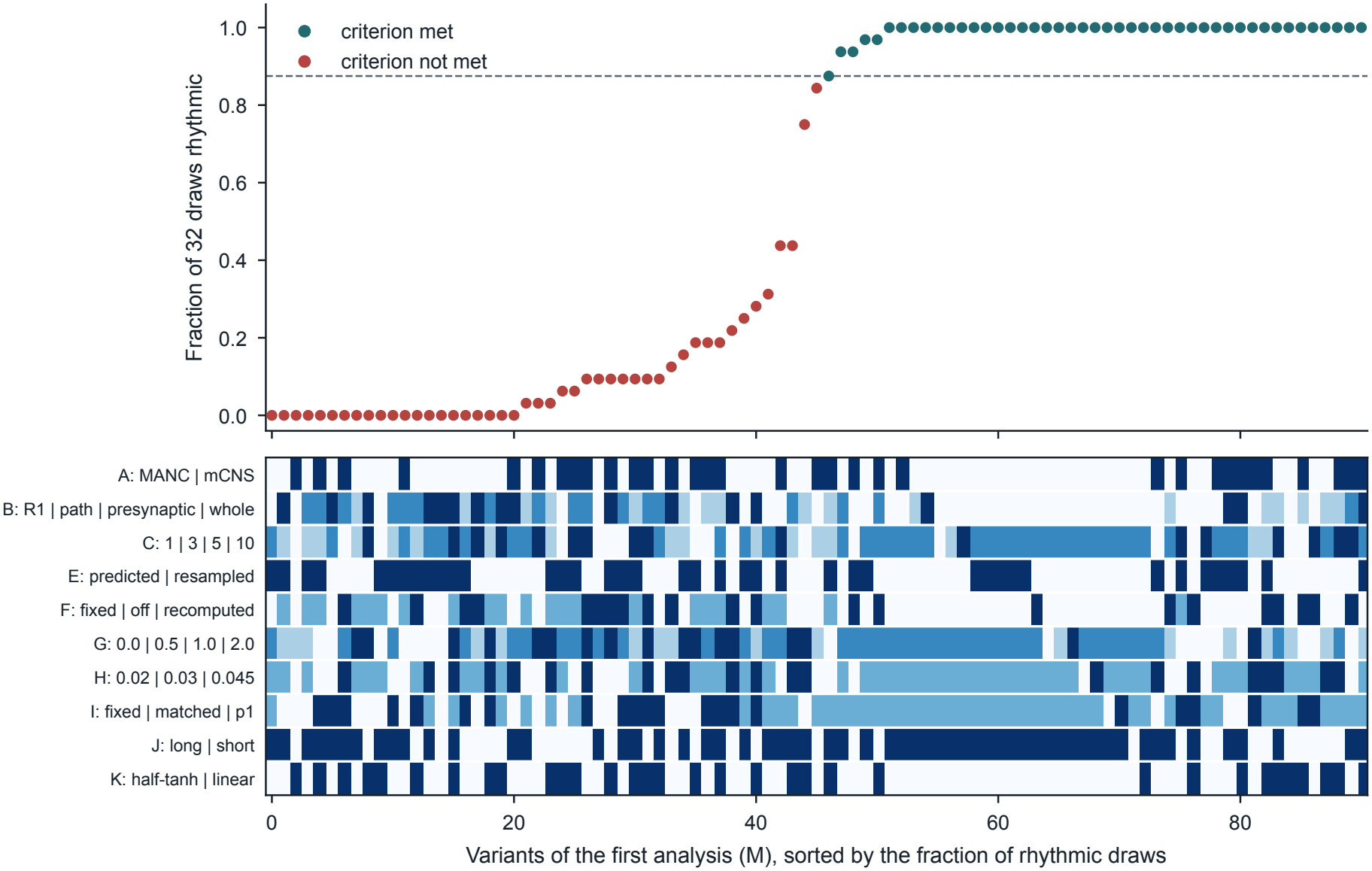
Specification curve for K1 in M, after DEVIATIONS part 5. Each column is one variant (U0, OAT and joint), sorted by the fraction of 32 draws with a rhythmic circuit score; the dashed line is the 28/32 threshold. Rows below mark each variant’s level in each dimension, from light (first listed) to dark. Several M levels were later found to change the model by accident; the label shown is M’s frozen label, not M2’s.

### S2 Further results of the robustness analysis

#### Rhythm at the assigned drive, by level

The proportion of the 48 size-normalized joint variants with a rhythm at the assigned drive differed by level, but each cell holds 9–24 variants and none enters a verdict: heterogeneity 0, 0.5, 1, 2 and 3 gave 4/10, 7/9, 7/11, 1/9 and 1/9; the recruitment-matched rule, the recruitment-band rule and the authors’ auto-tuning rule gave 12/17, 6/16 and 2/15; synaptic scale 0.02, 0.03 and 0.045 gave 1/16, 12/17 and 7/15; and the front-leg network, its expansion by path cells, the presynaptic expansion and the whole cord gave 6/12, 9/12, 3/12 and 2/12. In the test of the drive–frequency relation, a flag for a drive whose median count of active neurons in the default front-leg set exceeded the highest such count at the assigned drive marked the two highest drives in all 40 evaluable variants (and the next drive in 14), so it did not separate variants that met the criterion from those that did not.

#### Predictions

Nine of the plan’s ten predictions were true. The false one, that rhythm generation fails at threefold heterogeneity as a single change, depends on the lower bound on the time constant (Methods): it is false on the run as made (29/32), cannot be scored without the bound, and is true if those variants are treated as failed. The plan had also given a probability of 0.35 to a rhythm in at least 50% of the joint variants with size normalization on; the outcome was 20 of 48.

#### E2 homologues in the expanded networks

In the post hoc knockout check of the first analysis, silencing all left E2 homologues abolished the rhythm less often than silencing the T1 cell alone (0.55against 0.65 in the whole cord), and the remaining rhythm was slower (8.2–8.7 against 9.7–11.5 Hz before silencing). The second analysis pre-specified one contrast on this pattern, reporting numbers only, with no rule for interpreting them. It used one whole-cord network in which 31 of 32 draws were rhythmic before silencing (median 11.5 Hz). Silencing the T1 E2 cell abolished the rhythm in 20 of 31 draws, and the draws that stayed rhythmic had a median of 8.1 Hz. Silencing the left T1–T3 E2 homologues abolished it in 17 of 31 (8.7 Hz). Silencing the T1 E2 cell together with the left T2 and T3 homologues of E1, E2 and I1 abolished it in 29 of 31 (7.3 Hz in the two that stayed). We do not interpret these numbers.

**Table S1:**
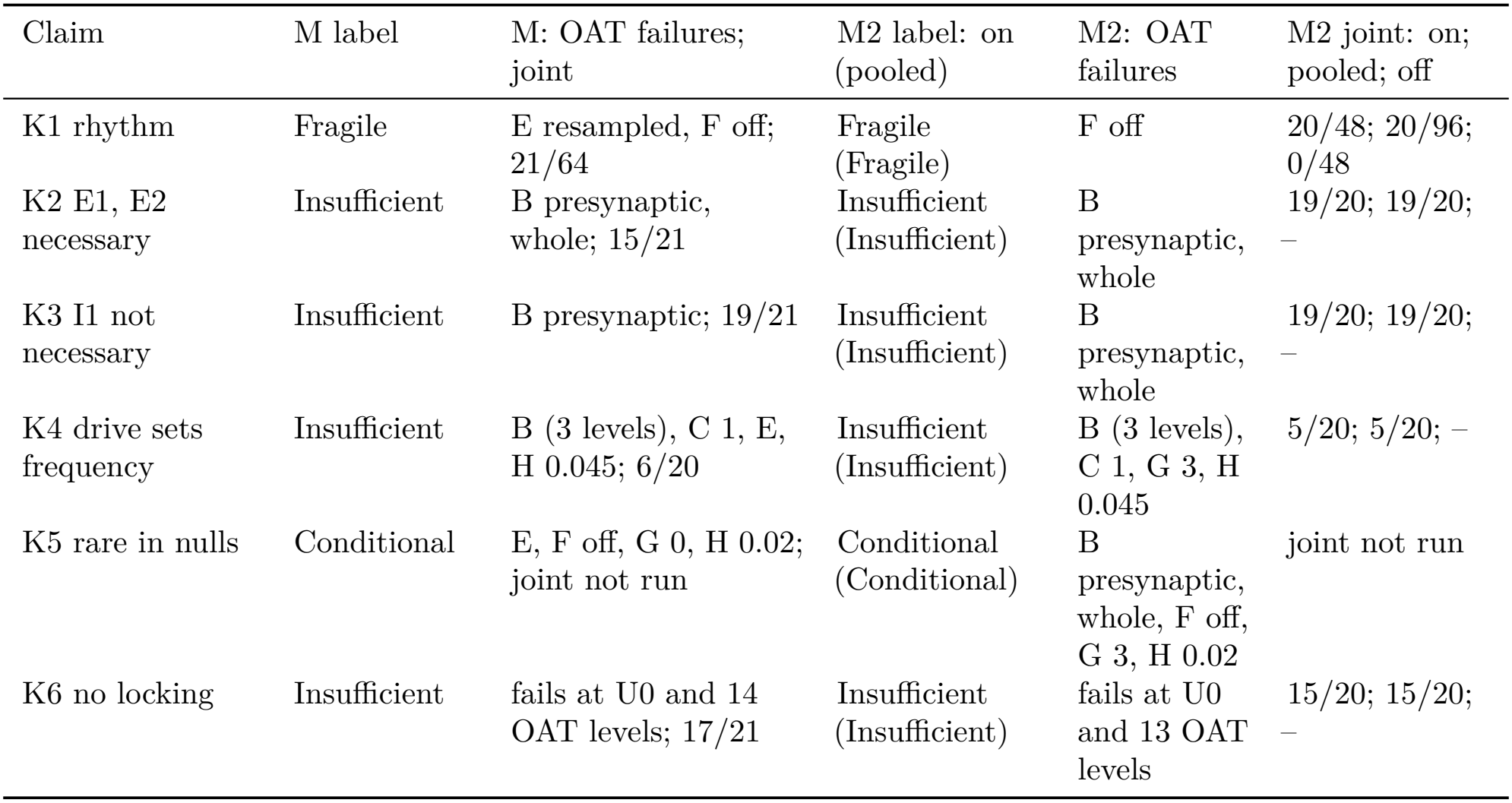
M and M2 side by side. M labels are frozen and include DEVIATIONS parts 1–5. Each M2 label for size normalization on is printed with the pooled label in the same cell; that denominator was chosen after M’s outcomes were read. Joint entries are passes over evaluable variants. M2 has 48 joint variants with size normalization on, 96 pooled and 48 off, and a label needs 75% of its denominator evaluable. K2–K4 and K6 were evaluable in no off variant, because K1 failed in all of them; K5 was not run in joint variants.

| Claim | M label | M: OAT failures; joint | M2 label: on (pooled) | M2: OAT failures | M2 joint: on; pooled; off |
|---|---|---|---|---|---|
| K1 rhythm | Fragile | E resampled, F off; 21/64 | Fragile (Fragile) | F off | 20/48; 20/96; 0/48 |
| K2 E1, E2 necessary | Insufficient | B presynaptic, whole; 15/21 | Insufficient (Insufficient) | B presynaptic, whole | 19/20; 19/20; – |
| K3 I1 not necessary | Insufficient | B presynaptic; 19/21 | Insufficient (Insufficient) | B presynaptic, whole | 19/20; 19/20; – |
| K4 drive sets frequency | Insufficient | B (3 levels), C 1, E, H 0.045; 6/20 | Insufficient (Insufficient) | B (3 levels), C 1, G 3, H 0.045 | 5/20; 5/20; – |
| K5 rare in nulls | Conditional | E, F off, G 0, H 0.02; joint not run | Conditional (Conditional) | B presynaptic, whole, F off, G 3, H 0.02 | joint not run |
| K6 no locking | Insufficient | fails at U0 and 14 OAT levels; 17/21 | Insufficient (Insufficient) | fails at U0 and 13 OAT levels | 15/20; 15/20; – |

### S3 Further results of the drive sweeps

#### Single-change variants

In the default model and the 20 single-change variants a rhythm was present in the sweep in 19. The whole-cord variant, which had a rhythm at its per-draw assigned currents (31 of 32 draws), reached at most 26 of 32 at a common drive of the sweep, and the variant without size normalization had at most 4 of 32 at its one feasible drive. At the best drive, silencing the E1 and the E2 homologues abolished the rhythm in 18 of 19 variants; the exception was the presynaptic expansion, where silencing E2 abolished it in 0.84 of draws. Silencing the I1 homologues left the rhythm in place in 17 of 19; the exceptions were the presynaptic expansion (abolished in 0.50 of draws) and threefold heterogeneity (0.14). Three variants had four or more usable drives, and *ρ* was negative in all three.

#### Refinement of the sweep

The refinement covered the 16 size-normalized joint variants with a rhythm neither in the coarse sweep nor at the assigned drive, and added 6–27 eighth-octave drives to each, between the highest coarse drive with fewer than five active neurons and the lowest with more than 500. It found a rhythm in two whole-cord variants with homogeneous parameters and the highest synaptic scale, in 32 of 32 draws; neither had any rhythmic draw at a feasible drive of the coarse sweep.

**Table S2:** M at U0 and at every OAT level (after DEVIATIONS part 5). Values from`multiverse/summary_M.json`. K2 gives E1 and E2 loss, K3 loss, K5 the difference and *p*, K6 the locked fraction.— means unevaluable because K1 failed. The E level is pooled over 8 resamples; its K1 entry gives the number of resamples that held.

| Variant | K1 (/32) | K2 | K3 | K4 *ρ* | K5 | K6 |
|---|---|---|---|---|---|---|
| U0 | 32 | 1.00, 0.97 | 0.00 | 1.00 | 0.963, 0.020 | 0.312 |
| A: male CNS (VNC ROIs) | 32 | 1.00, 1.00 | 0.00 | 0.90 | 0.801, 0.030 | 0.344 |
| B: path | 32 | 1.00, 0.97 | 0.00 | 0.70 | 0.961, 0.030 | 0.344 |
| B: presynaptic | 30 | 0.40, 0.80 | 0.17 | −0.90 | 0.500, 0.030 | 0.094 |
| B: whole | 32 | 0.28, 0.84 | 0.06 | −0.80 | 0.750, 0.030 | 0.094 |
| C: 1 | 32 | 1.00, 1.00 | 0.00 | 0.70 | 0.992, 0.030 | 0.438 |
| C: 3 | 32 | 1.00, 1.00 | 0.00 | 0.90 | 0.980, 0.030 | 0.312 |
| C: 10 | 32 | 1.00, 1.00 | 0.00 | 0.90 | 0.961, 0.030 | 0.375 |
| E: resampled (8) | 6/8 held | — | — | 2/6 held | 6/8 held | — |
| F: recomputed | 32 | 1.00, 0.97 | 0.00 | 1.00 | 0.977, 0.030 | 0.312 |
| F: off | 10 | — | — | — | 0.266, 0.152 | — |
| G: 0.0 | 32 | 1.00, 1.00 | 0.00 | 1.00 | 0.969, 0.061 | 0.312 |
| G: 0.5 | 32 | 1.00, 1.00 | 0.00 | 1.00 | 0.961, 0.030 | 0.281 |
| G: 2.0 | 32 | 1.00, 0.97 | 0.00 | 0.90 | 0.852, 0.030 | 0.156 |
| H: 0.02 | 32 | 1.00, 1.00 | 0.00 | 0.90 | −0.023, 1.000 | 0.312 |
| H: 0.045 | 32 | 1.00, 0.94 | 0.00 | 0.00 | 0.965, 0.030 | 0.344 |
| I: fixed 250 | 32 | 1.00, 0.97 | 0.00 | 1.00 | 0.926, 0.030 | 0.312 |
| I: P1 tuner | 32 | 1.00, 0.97 | 0.00 | 1.00 | 0.977, 0.030 | 0.312 |
| J: long | 32 | 1.00, 0.97 | 0.00 | 1.00 | 0.984, 0.030 | 0.312 |
| K: threshold-linear | 32 | 1.00, 0.97 | 0.00 | 1.00 | 0.949, 0.030 | 0.312 |

In the 13 refined variants with two- or threefold heterogeneity the largest number of rhythmic draws at a common feasible drive rose by at most 6 and reached at most 27. Knockouts were not repeated at refined drives. With the drive chosen draw by draw over both sweeps, 8 of the 16 reached 28 rhythmic draws. One further variant, a presynaptic-expansion variant with default heterogeneity that had a rhythm at its assigned drive, was not refined and has 14 such draws in the coarse sweep.

### S4 Additional table

**Table S3:** Key R1 interventions. Frequency shifts are summarized only when more than one paired draw had rhythmic motor neurons in both conditions.

| Condition | Draws | Fraction rhythmic | Score | Frequency shift (Hz) |
|---|---|---|---|---|
| Left DNg100, *I* = 250 | 64 | 1.000 | 0.957 | — |
| Minimal circuit | 64 | 0.984 | 0.985 | — |
| KO E1 | 16 | 0.000 | 0.000 | NA |
| KO E2 | 16 | 0.062 | 0.058 | NA (*n* = 1) |
| KO I1 | 16 | 1.000 | 0.878 | −1.15 |

**Figure S2:**
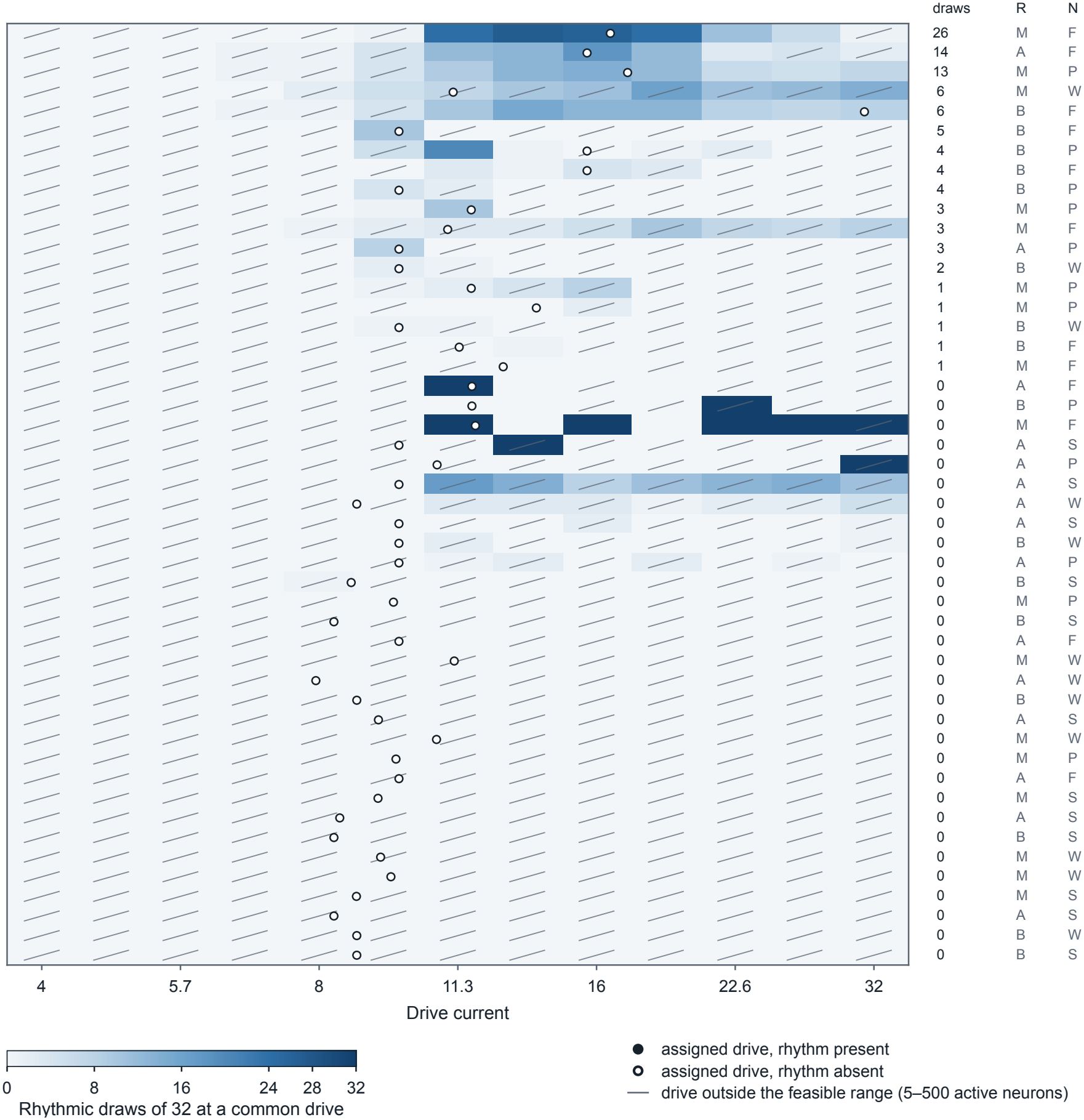
Rhythmic draws across drive in the 48 joint variants without size normalization. Conventions as in Fig. 4; drives run from 4 to 32 in quarter-octave steps.

